# Cryo-EM reveals a new allosteric binding site at the M_5_ mAChR

**DOI:** 10.1101/2025.02.05.636602

**Authors:** Wessel A. C. Burger, Jesse I. Mobbs, Bhavika Rana, Jinan Wang, Keya Joshi, Patrick R. Gentry, Mahmuda Yeasmin, Hariprasad Venugopal, Aaron M. Bender, Craig W. Lindsley, Yinglong Miao, Arthur Christopoulos, Celine Valant, David M. Thal

## Abstract

The M_5_ muscarinic acetylcholine receptor (M_5_ mAChR) represents a promising therapeutic target for neurological disorders. However, the high conservation of its orthosteric binding site has posed significant challenges for drug development. While selective positive allosteric modulators (PAMs) offer a potential solution, a structural understanding of the M_5_ mAChR and its allosteric binding sites has remained limited. Here, we present a 2.8 Å cryo-electron microscopy structure of the M_5_ mAChR complexed with heterotrimeric G_q_ protein and the agonist iperoxo, completing the active-state structural characterization of the mAChR family. To identify the binding site of M_5_-selective PAMs, we implemented an integrated approach combining mutagenesis, pharmacological assays, structural biology, and molecular dynamics simulations. Our mutagenesis studies revealed that selective M_5_ PAMs bind outside previously characterized M_5_ mAChR allosteric sites. Subsequently, we obtained a 2.1 Å structure of M_5_ mAChR co-bound with acetylcholine and the selective PAM VU6007678, revealing a novel allosteric pocket at the extrahelical interface between transmembrane domains 3 and 4 that was confirmed through mutagenesis and simulations. These findings demonstrate the diverse mechanisms of allosteric regulation in mAChRs and highlight the value of integrating pharmacological and structural approaches to identify novel allosteric binding sites.

## Introduction

The M_5_ muscarinic acetylcholine receptor (mAChR) belongs to the class A G protein-coupled receptor (GPCR) family. As one of five mAChR subtypes (M_1_-M_5_), it responds to the endogenous neurotransmitter acetylcholine (ACh)^1^. Despite its low expression levels in the central nervous system (CNS), the M_5_ mAChR plays crucial roles, primarily localizing to dopamine-containing neurons in the substantia pars nigra and ventral tegmental areas^2–4^. Historically, drug discovery efforts targeting the M_5_ mAChR concentrated on antagonists and negative allosteric modulators (NAMs). This focus stemmed from evidence that M_5_ mAChR inactivation can modulate dopaminergic signalling in the CNS^5,6^, suggesting therapeutic potential for substance addiction, depression, and anxiety^7–15^. However, M_5_ mAChR activation may offer equally promising therapeutic applications. Studies using M_5_ mAChR knockout mice revealed that this receptor mediates CNS vasculature dilation, thereby regulating cerebral blood flow^16,17^. This finding suggests that selective M_5_ mAChR activation could benefit conditions such as Alzheimer’s disease, schizophrenia, and ischemic stroke by enhancing CNS circulation and blood flow.

The development of selective M_5_ mAChR activators faces significant challenges, primarily due to the highly conserved orthosteric acetylcholine binding site shared across all five mAChR subtypes^18,19^. This conservation has historically hindered the development of subtype-selective orthosteric ligands. To overcome this limitation, researchers shifted focus to allosteric modulators, which target spatially distinct binding sites^20^. This strategy proved promising, with several selective allosteric modulators successfully developed for the M_5_ mAChR^21–26^. The selectivity of these modulators likely stems from non-conserved residues present in the allosteric binding sites, distinguishing them from the highly conserved orthosteric site.

M_5_-selective positive allosteric modulators (PAMs) were initially discovered and developed based on a bromo isatin scaffold, as exemplified by the PAM VU0119498^27^. VU0119498 enhanced ACh signalling in a calcium assay at all Gα_q_ coupled mAChRs (M_1_, M_3_, M_5_) and was therefore chosen as a tool compound for further M_5_ mAChR PAM optimization. A range of new M_5_ mAChR PAMs were developed based on VU0119498 and these all show M_5_ mAChR selectivity over all mAChR subtypes with varying levels of potency and PAM activity^21,22,24^. Unfortunately, all M_5_ mAChR selective PAMs based on the isatin scaffold exhibit a poor drug metabolism and pharmacokinetic (DMPK) profile that was hard to overcome due to the intractability of the isatin scaffold in medicinal chemistry optimization^1,24^.

To overcome these limitations, high-throughput screening identified a novel scaffold amenable to modification^25^. This effort led to the development of 1-((1*H*-indazol-5-yl)sulfoneyl)-*N*-ethyl-*N*-(2-(trifluoromethyl)benzyl)piperidine-4-carboxamide (ML380), which emerged as the most potent M_5_-selective PAM at the time^25^. ML380’s ability to penetrate the CNS made it valuable for studying molecular mechanisms of allosteric modulation and as a template for further derivatives^26,28,29^. However, poor partition coefficients ultimately limited its *in vivo* utility^25^. Despite these incremental advances in developing M_5_-selective PAMs, their precise allosteric binding site remained unknown, significantly limiting rational, structure-based drug design. The M_5_ mAChR contains at least three known allosteric sites: the ‘mAChR common’ extracellular vestibule (ECV), the amiodarone-binding site^30^, and the extrahelical EH4 pocket recognized by the M_5_-selective NAM ML375^31^. However, none of these sites were structurally confirmed as the binding location for M_5_-selective PAMs. Understanding the location of M_5_- selective PAM binding site will accelerate the development of improved compounds for *in vivo* use^32^. Therefore, we initiated studies to determine the binding site of M_5_-selective PAMs, beginning with ML380.

## Results

### ML380 does not bind to known M_5_ mAChR allosteric binding sites

To identify ML380’s binding site, we first investigated its interaction with known allosteric sites through functional inositol monophosphate (IP1) accumulation assays. We tested ML380’s activity on both wild-type (WT) M_5_ mAChR and mutants targeting two known allosteric sites: (1) the ‘common’ extracellular vestibule (ECV) allosteric site^33,34^, using alanine mutants, and (2) the EH4 pocket used by the M_5_-selective NAM ML375^31^, using mutations that convert non-conserved residues to their M_2_ mAChR equivalents (A113^3.35^V, G152^4.47^A, and L156^4.51^V). These experiments measured three key parameters: ML380’s affinity (p*K*_B_), its efficacy in the system (log τ), and its functional cooperativity with ACh (log αβ)^35^. Consistent with previous studies, ML380 demonstrated both agonism and positive cooperativity with ACh at the WT M_5_ mAChR^28,29^ (Fig. 1a). The EH4 pocket mutant maintained ML380 function with no significant change in affinity compared to WT (Fig. 1a,c,d, Table 1). While some ECV alanine mutants showed significant differences in ML380’s affinity, efficacy, and cooperativity parameters, the compound retained its ability to bind, activate, and modulate receptor function in all cases (Fig. 1b,c-e, Supplementary Fig. 1, Supplementary Table 1). These results indicated that ML380 binds neither to the common ECV allosteric site nor to the EH4 pocket used by ML375, leading us to use cryo-electron microscopy (cryo-EM) to identify its binding site.

**Figure 1.**
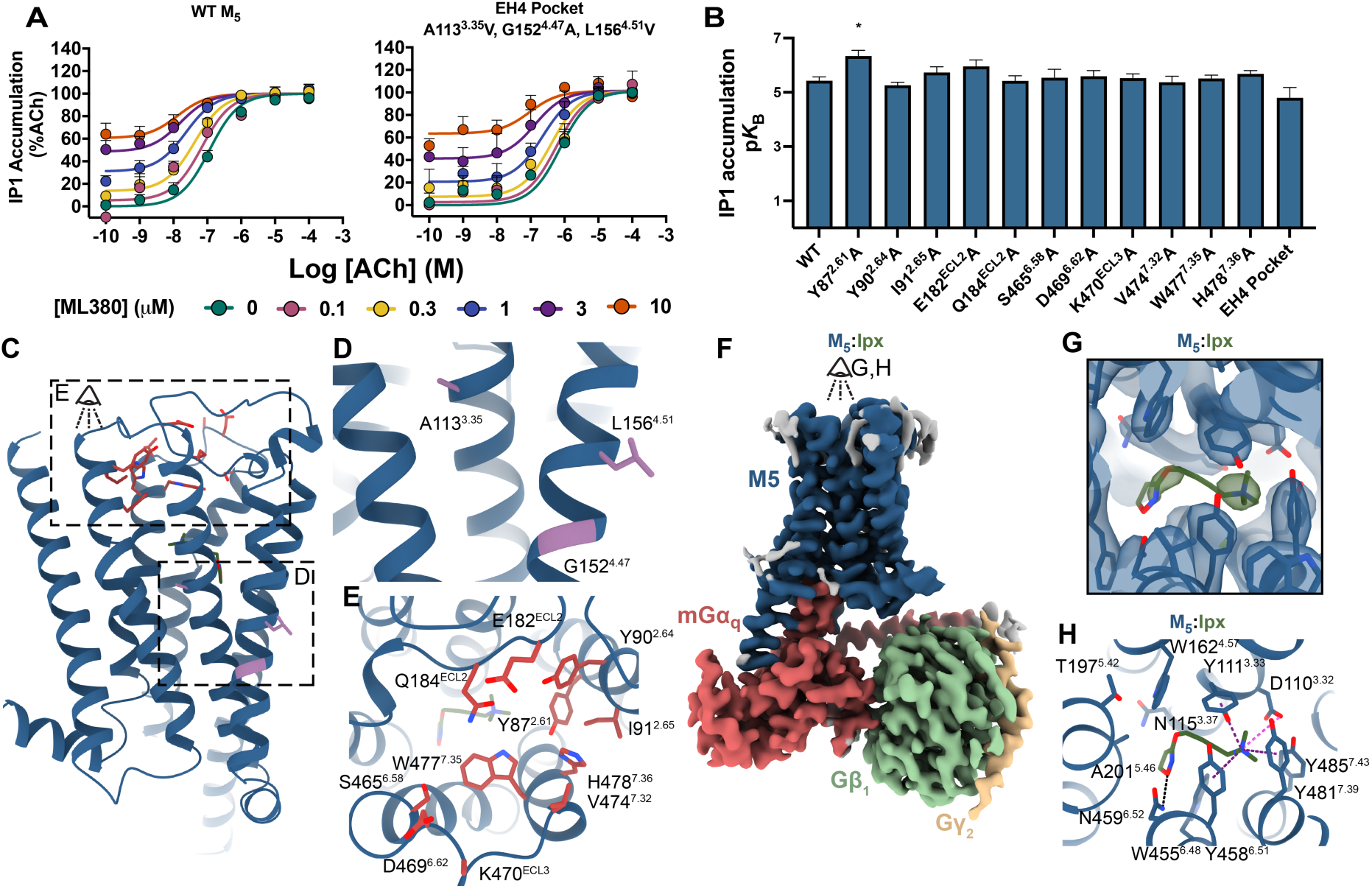
Structural and functional analysis of ligand-receptor interactions at the M_5_ mAChR. **(A)** Interaction of ML380 with ACh in an IP1accumulation assay at WT M_5_ mAChR (left) or M_5_ EH4 pocket mutant mAChR (right) expressing CHO cells. Data points represent mean ± SEM of three to seven individual experiments performed in duplicate. WT M_5_ mAChR *n* = 7, M_5_ EH4 pocket mutant *n =* 3. An operational model of allosterism was fit to the data. **(B)** Effects of the M_5_ mAChR mutations on the p*K*_B_ of ML380. Data represent the mean ± SEM of three to seven independent experiments performed in duplicate. WT M_5_ mAChR *n* = 7, all other mutants *n =* 3. *, significantly different from WT, p < 0.05, one-way ANOVA, Dunnett’s post hoc test. Parameters obtained are listed in Supplementary Table 1. **(C)** M_5_ mAChR mutated residues are shown on the structure of the M_5_ mAChR. **(D)** Residues of the EH4 pocket are shown as purple sticks. **(E)** Residues of the ‘common’ ECV are shown as red sticks. **(F)** Consensus cryo-EM map of the M_5_ mAChR in complex with Gα_mGsQi_/Gβ_1_γ_2_ bound to iperoxo resolved to 2.8 Å (FSC 0.143). The receptor is shown in dark blue, the heterotrimeric G_q_ protein is shown in red, green, and yellow for the α, β, γ subunits, respectively. **(G)** Cryo-EM density (local refined receptor map, contour level 0.36) for iperoxo in the orthosteric binding site. **(H)** Interactions of iperoxo with the orthosteric binding site. Charge-charge interactions are shown as pink dotted lines, hydrogen bonds are shown as black dotted lines while purple dotted lines represent a cation-π interaction with Y111^3.33^, W455^6.48^, Y458^6.51^, and Y481^7.39^.

### Cryo-electron microscopy structure determination

Active state structures of the M_1_-M_4_ mAChRs have been determined by cryo-EM^34,36,37^, leaving the M_5_ mAChR as the last remaining subtype. To determine an active state M_5_ mAChR structure, we engineered a modified receptor construct with several key alterations: removal of intracellular loop 3 (ICL3) residues 237-421, addition of an N-terminal HA signal sequence, and incorporation of an anti-Flag epitope tag. This modified receptor was then fused to mini-Gα_qiN_ yielding a chimeric construct^38,39^. The receptor-G protein complex was prepared through a multi-step process including detergent solubilization for purification, stabilization with scFv16^40^, addition of apyrase to hydrolyze GDP, and inclusion of 10 µM iperoxo. Prior to cryo-EM analysis, we incubated the iperoxo-bound M_5_ mAChR-mini-Gα_qiN_ complex with 10 µM ML380 overnight on ice, followed by freezing and imaging using single-particle cryo-transmission electron microscopy (TEM) on a Titan Krios microscope. The resulting structure achieved a global resolution of 2.8 Å, providing sufficient EM density maps to position the receptor and Gα_qiN_β_1_γ_2_ complex (Fig. 1f, Supplementary Fig. 2-3, Supplementary Table 2). However, due to poor density, the scFv16 component was excluded from subsequent data analysis and modelling.

### Analysis of the active state M_5_ mAChR

The iperoxo-bound M_5_ mAChR-mini-Gα_qiN_ complex revealed iperoxo bound to the canonical mAChR orthosteric binding site, which is characterized by several key aromatic residues (Fig. 1g-h). The rotamer toggle switch residue W455^6.48^ forms the binding site’s floor, while three tyrosine residues (Y111^3.33^, Y458^6.51^, and Y481^7.39^) move inward during activation to create a tyrosine lid, separating the orthosteric site from the ECV. Consistent with previous M_1_-M_4_ mAChR structures^33,34,36,37^, iperoxo was positioned between these aromatic residues (Fig. 1h). Additional binding site residues include the aromatic residues W162^4.57^ and Y485^7.43^, along with non-aromatic residues D110^3.32^, N115^3.37^, T194^5.39^, T197^5.42^, A201^5.46^, N459^6.52^, L188^ECL2^, and C484^7.42^. Two residues play crucial roles in iperoxo recognition. Residue N459^6.52^ forms hydrogen bonds with iperoxo, while D110^3.32^ engages in a charge-charge interaction with iperoxo’s quaternary nitrogen.

The active state M_5_ mAChR displays all the hallmarks of an active state class A GPCR. Relative to the inactive state, tiotropium bound M_5_ mAChR, there is an 8.1 Å outward movement of transmembrane 6 (TM6) and 4.4 Å inward movement of TM7 (Supplementary Fig. 4). At the extracellular face, there is an outward movement of TM5 and inward movement of TM6 and extracellular loop 3 (ECL3) that lead to a contraction of the ECV (Supplementary Fig. 4b). These global changes in TM and ECL movements are mediated by changes in the conserved class A activation motifs including the D^3.49^R^3.50^Y^3.51^, P^5.50^V^3.40^F^6.44^, and N^7.49^P^7.50^xxY^7.53^ motifs (Supplementary Fig. 4D, superscript refers to the Ballesteros and Weinstein scheme for conserved class A GPCR residues)^41,42^.

Our active iperoxo bound M_5_ mAChR structure enables the first comprehensive analysis of the entire mAChR family in their active state. Comparing this M_5_ mAChR structure with other iperoxo-bound mAChR structures reveals remarkable similarity, with root mean squared deviation (RMSD) values of 0.49-0.77 Å (Fig. 2a) ^33,34,36,37^. The TM domains, ECL domains, orthosteric site residues, and iperoxo positioning are nearly identical across all mAChR subtypes (Fig. 2b-d). At the receptor-G protein interface, Gα_q_- and Gα_i/o_-coupled mAChRs show distinct α5 helix insertion angles into the TM bundle (Fig. 2f-g). In Gα_q_-coupled receptors (M_1_, M_3_, and M_5_ mAChRs), the α5 helix top rotates toward TM6, while in Gα_i/o_-coupled receptors (M_2_ and M_4_ mAChRs), it rotates toward TM2. This pattern, however, does not extend to the G protein N-terminus (Fig. 2e), possibly due to the varying use of scFv16, Nb35, and N-terminal chimeras across different structures. Additionally, the Gα_q_-coupled M_1_, M_3_, and M_5_ mAChR structures display a more extended and resolved TM5.

**Figure 2:**
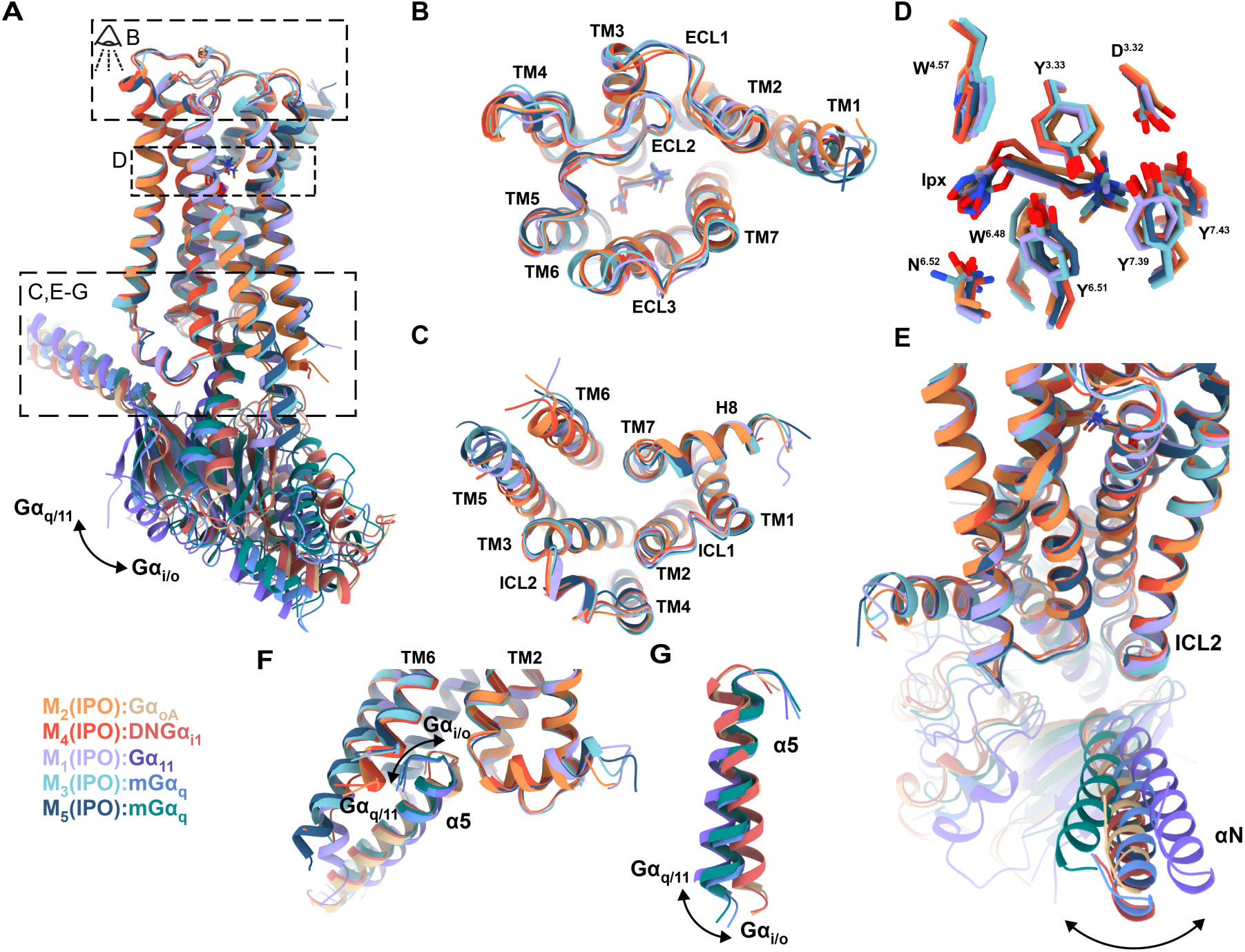
Structural comparison of iperoxo bound, active state M_1_-M_5_ mAChR structures determined by cryo-EM. **(A)** Overall view of the M_1_ to M_5_ mAChRs complexed to Gα and bound to iperoxo. M_1_-iperoxo-Gα_11_ is coloured purple, M_2_-iperoxo-Gα_oA_ is coloured orange, M_3_-iperoxo-Gα_mGsQi_ is coloured light blue, M_4_-iperoxo-dnGα_i1_ is coloured red, M_5_-iperoxo-Gα_mGsQi_ is coloured dark blue. **(B)** Extracellular view comparing ECLs and TM regions across the M_1_ to M_5_ mAChRs. **(C)** Intracellular view (G protein removed) comparing ICLs and TM regions across the M_1_ to M_5_ mAChRs. **(D)** Overlay of iperoxo and orthosteric binding site residues at the M_1_ to M_5_ mAChRs. **(E)** Intracellular view comparing the αN movement of Gα at M_1_ to M_5_ mAChRs. Changes in orientation is indicated by an arrow. **(F)** Intracellular view comparing the α5 insertion of Gα into the M_1_ to M_5_ mAChRs. Changes in insertion is indicated by an arrow. **(G)** Intracellular view comparing the α5 rotation of Gα relative to M_1_ to M_5_ mAChRs. Changes in rotation is indicated by an arrow.

In line with our mutagenesis results (Fig. 1a-b), no cryo-EM density was observed for ML380 in the ‘common’ ECV allosteric site or in the EH4 binding pocket (Fig. 3a-b). Following focused refinement with a mask, some cryo-EM density was observed parallel to TM1 and TM7 in the ECV (coloured green Fig. 3c), however, this density was commonly observed in other mAChR structures and is likely a lipid molecule^43^. We also observed partial density directly below the EH4 pocket at the bottom of the TM2,3,4 interface (coloured orange in Fig. 3c). To investigate whether this partial density reflects ML380 occupancy at its allosteric binding site, we conducted radioligand binding experiments using M_5_/M_2_ TM chimera mutants^31^. We measured interactions between ACh and [^3^H]-*N*-methyl scopolamine ([^3^H]-NMS) with increasing ML380 concentrations. If ML380 binds at the TM2,3,4 interface, we would expect complete loss of its allosteric modulation in both M_5_/M_2_ TM2,3,4 and M_5_/M_2_ TM3,4,5 chimeras, given ML380’s selectivity profile^25^. Indeed, swapping TM2-5 completely abolished ML380’s ability to modulate ACh binding (Fig. 3d, Supplementary Table 3). The M_5_/M_2_ TM1,7,h8 chimera showed increased ML380 cooperativity, confirming that the density parallel to TM1 and TM7 is not ML380, while suggesting that exchanging these TMs affects the receptor’s global activation dynamics (Fig. 3d, Supplementary Table 3).

**Figure 3:**
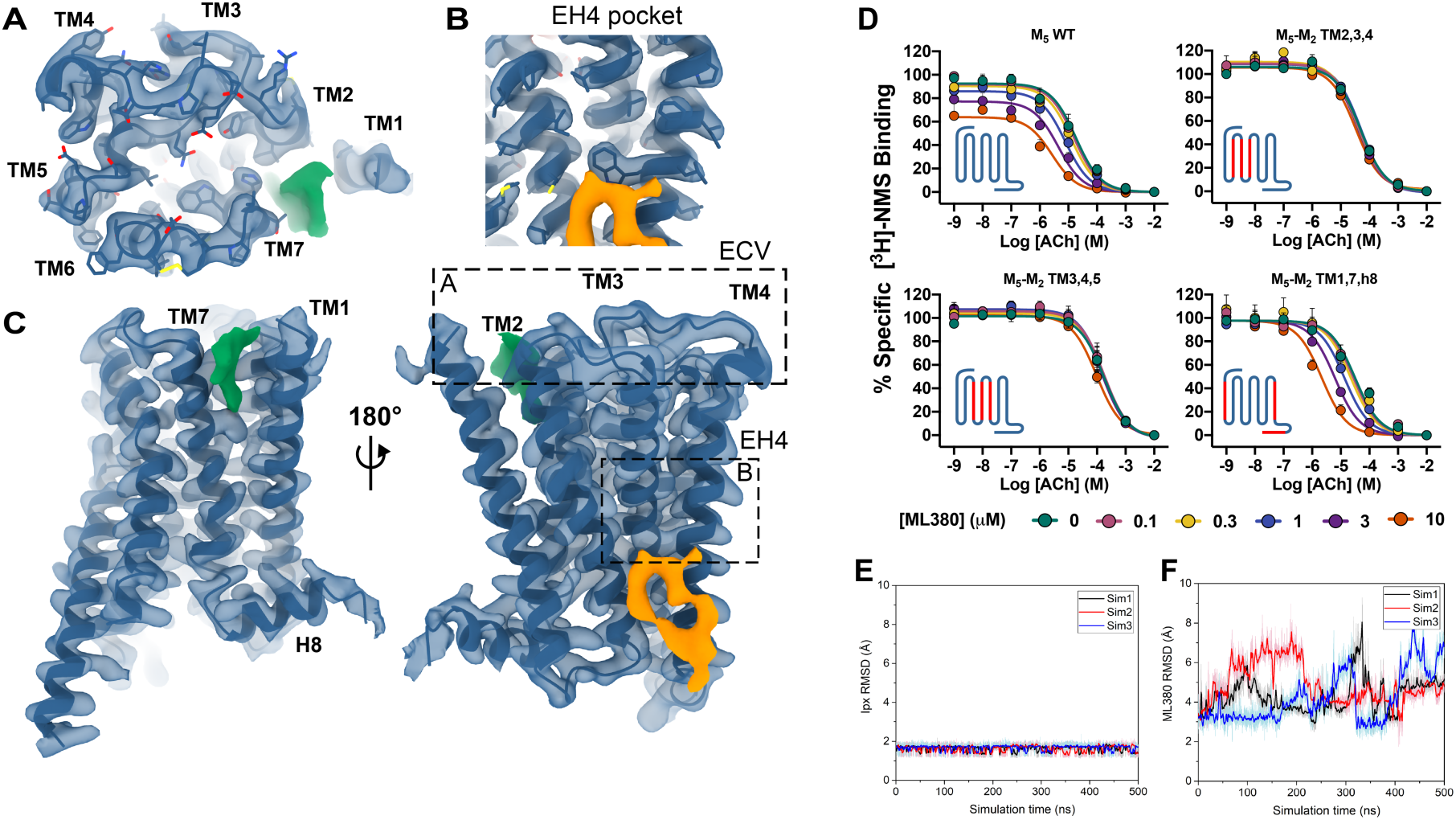
Potential cryo-EM density for ML380. **(A-C)** Local refined receptor cryo-EM map of the M_5_ mAChR (contour level 0.3). No density for ML380 was observed in the (A) ‘common’ ECV (B) or in the EH4 pocket. (C) potential density for ML380 was observed parallel to TM1 and TM7 (coloured green) and at the bottom of the TM2,3,4 interface (coloured orange). **(D)** [^3^H]-NMS equilibrium radioligand binding studies between [^3^H]-NMS, ACh and ML380 at the WT M_5_ mAChR and M_5_-M_2_ TM chimeric swaps. The insets are cartoons of the WT M_5_ mAChR and M_5_-M_2_ TM chimeric swaps with blue representing M_5_ mAChR domains and red representing M_2_ mAChR domains. Data points represent mean ± SEM of three individual experiments performed in duplicate. Data was fit to an allosteric ternary complex model. Parameters obtained are listed in Supplementary Table 3. **(E-F)** Root mean square deviations (RMSDs) of (E) iperoxo in the orthosteric binding pocket and (F) ML380 at the potential allosteric binding site at the bottom of the TM2,3,4 interface calculated from Gaussian accelerated molecular dynamics (GaMD) simulations of the cryo-EM structure, each performed with three separate replicates indicated with different coloured traces.

Although the TMs 2-5 mutagenesis results were promising, the density at the TM2,3,4 interface coincides with a common cholesterol binding site in class A GPCRs^44,45^, suggesting it might represent cholesterol or another lipid. For further analysis, we performed all-atom Gaussian accelerated Molecular Dynamics (GaMD) simulations on the modelled structure of the iperoxo-M_5_ mAChR-mini-Gα_qiN_ complex with ML380 bound at the TM2,3,4 interface (Fig. 3e-f). While iperoxo maintained its cryo-EM conformation with RMSD values mostly below 2 Å (Fig. 3e), ML380 showed significant fluctuations with RMSD values of ∼3-8 Å compared to the initial cryo-EM pose (Fig. 3f). The combination of mutagenesis data, GaMD simulations, and ambiguous cryo-EM density did not provide convincing support for assigning ML380 to this putative allosteric binding site.

### Use of an improved PAM, VU6007678, for structure determination

To discover the allosteric binding site for selective PAMs at the M_5_ mAChR, we investigated VU6007678^26^, an optimized derivative of ML380’s indanyl core. VU6007678 demonstrated enhanced M_5_ mAChR affinity, better positive cooperativity with ACh compared to ML380^29^, and significantly improved GPCR-G protein complex stabilization. To increase our chances of obtaining a PAM-bound complex, we implemented several biochemical and pharmacological modifications. We supplemented Nb35 with scFv16 during purification for enhanced complex stability, used the endogenous agonist ACh instead of iperoxo, and maintained VU6007678 at 10 µM throughout purification rather than adding it before grid freezing as with ML380. These modifications improved protein purification efficiency, yielding a sample at 18 mg/mL. When applied to Au grids, the sample produced a complex resolved to 2.1 Å from 418,794 particles (Fig. 4a, Supplementary Fig. 2-3).

**Figure 4:**
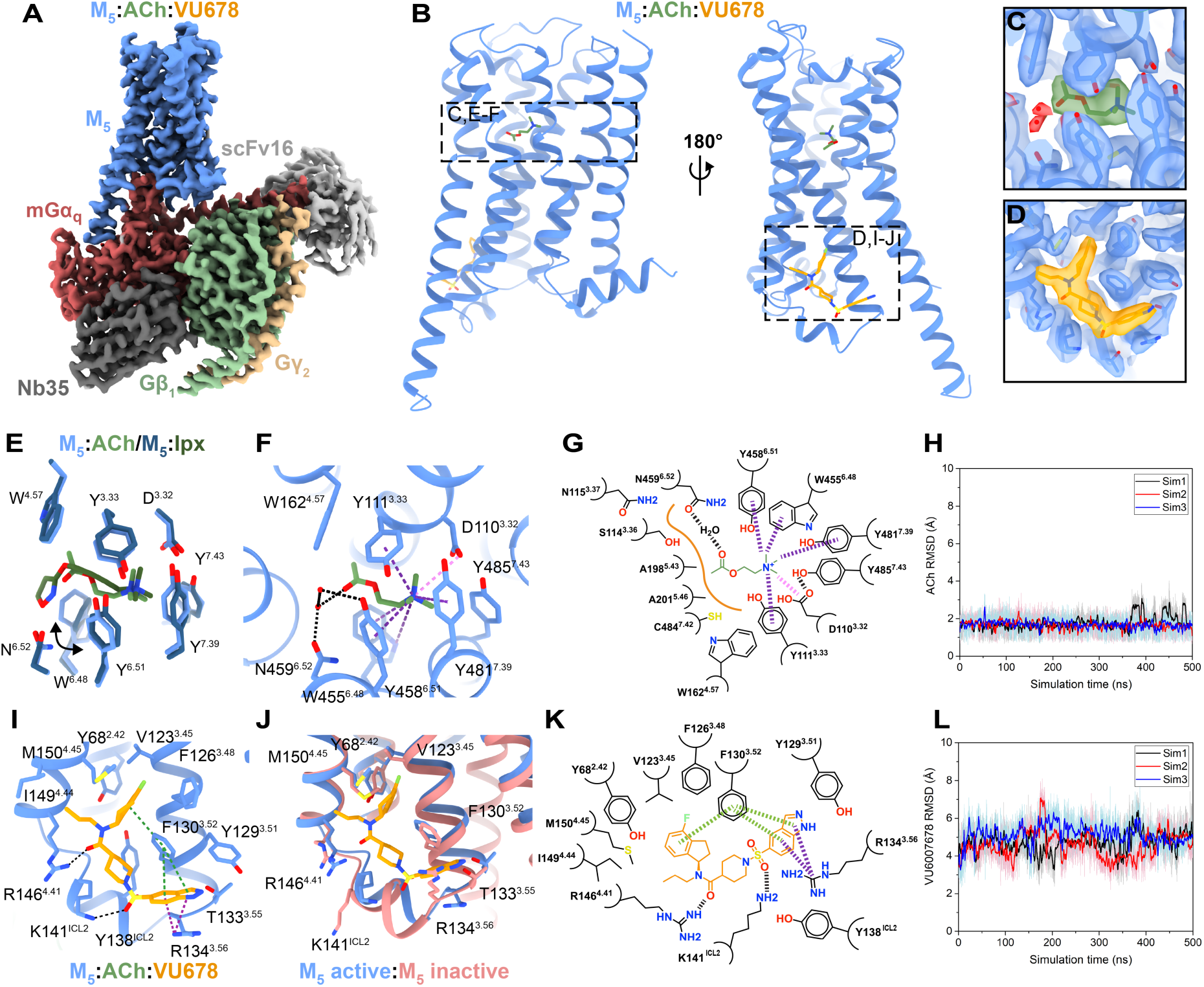
High resolution structure of ACh and VU6007678 bound M_5_ mAChR in complex with heterotrimeric G protein. **(A)** Consensus cryo-EM map of the M_5_ mAChR in complex with Gα_mGsQi_/Gβ_1_γ_2_ bound to ACh and VU6007678 resolved to 2.1 Å (FSC 0.143). The receptor is shown in light blue, the heterotrimeric G_q_ protein is shown in red, green, and yellow for the α, β, γ subunits, respectively, scFv16 is shown in light grey and Nb35 is shown in dark grey. **(B)** Model of the M_5_ mAChR showing ACh (green) in the orthosteric binding site and VU6007678 (orange) in an allosteric binding site at the bottom at TM3 and 4 and on top of ICL2. **(C)** Cryo-EM density (local refined receptor map, contour level 0.43) for iperoxo in the orthosteric binding site. **(D)** Cryo-EM density (local refined receptor map, contour level 0.43) for VU6007678 in an allosteric binding site. **(E)** Comparison of the orthosteric binding site interactions at ACh and Ipx-bound M_5_ mAChR. Shown in light blue sticks is ACh-bound M_5_ mAChR residues, dark blue sticks is Ipx-bound M_5_ mAChR residues. ACh is shown in light green sticks and Ipx is shown in dark green sticks. **(F)** Interactions of ACh with the orthosteric binding site of the M_5_ mAChR. Charge-charge interactions are shown as pink dotted lines, hydrogen bonds are shown as black dotted lines while purple dotted lines represent a cation-π interaction with Y111^3.33^, W455^6.48^, Y458^6.51^, and Y481^7.39^. Red spheres indicate water molecules. **(G)** 2D interaction plot of ACh with the orthosteric binding site of the M_5_ mAChR. Charge-charge interactions are shown as pink dashed lines, hydrogen bonds are shown as black dashed lines, cation-π interactions are shown as purple dashed lines and hydrophobic interactions are shown as a solid orange line. **(H)** RMSDs of ACh relative to the starting conformation in the orthosteric pocket calculated from GaMD simulations of the cryo-EM structure performed with three separate replicates indicated through different coloured traces. **(I)** Interactions of VU6007678 with its allosteric binding site at the M_5_ mAChR. Hydrogen bonds are shown as black dotted lines, while purple dotted lines represent cation-π interactions, and green dotted lines represent π-π interactions. **(J)** Comparison of the VU6007678 allosteric binding site to the inactive M_5_ mAChR when bound to the orthosteric antagonist tiotropium (PDB:6OL9). VU6007678 is shown in orange, the M_5_ mAChR bound to ACh and VU6007678 in light blue and the inactive tiotropium bound M_5_ mAChR in salmon. **(K)** 2D interaction plot of VU6007678 with the its allosteric binding site. Hydrogen bonds are shown as black dashed lines, while purple dashed lines represent cation-π interactions, and green dashed lines represent π-π interactions. **(L)** RMSDs of VU6007678 relative to the starting conformation at the allosteric binding site calculated from GaMD simulations of the cryo-EM structure performed with three separate replicates indicated through different coloured traces.

The high-quality cryo-EM density maps enabled precise placement of the receptor, Gα_qiN_ β_1_γ_2_, Nb35, and scFv16, with clear sidechain orientations for most amino acids. The ACh- VU6007678 structure closely resembles the iperoxo structure, showing an RMSD value of 0.49 Å (Supplementary Fig. 5a-d). The orthosteric binding pocket shows well-resolved density for ACh positioned beneath the tyrosine lid residues and above W455^6.48^ (Fig 4b-c). While ACh engages the same orthosteric binding pocket residues as iperoxo with similar orientations, W455^6.48^ adopts a more horizontal and planar orientation with ACh-bound, similar to observations in the M_4_ mAChR with ACh and iperoxo^34^ (Fig. 4e). ACh forms key interactions within the orthosteric binding pocket through its quaternary ammonium group and acetyl moiety (Fig. 4f,g). The positively charged quaternary ammonium establishes a cation-π interaction with the aromatic cage formed by Y111^3.33^, W455^6.48^, Y458^6.51^, and Y481^7.39^, while also engaging in a charge-charge interaction with D110^3.32^. The acetyl group of ACh forms a hydrogen bond with a water molecule that coordinates with N459^6.52^ (Fig. 4g) altogether anchoring ACh in an orientation that promotes receptor activation.

Clear, well-defined density unambiguously corresponding to VU6007678 was detected at the intracellular interface of TM3,4 and above ICL2 (Fig. 4a, b, d). The extended binding site accommodates VU6007678 through multiple interactions with the M_5_ mAChR (Fig. 4i, k). The indanyl core, positioned at the top of the allosteric binding site, forms hydrophobic interactions with M150^4.45^, F126^3.48^, and V123^3.45^, while engaging in an edge-to-face π-interaction with F130^3.52^. The propyl chain extends toward TM4, forming a hydrophobic interaction with I149^4.44^. Hydrogen bonding occurs between the carboxamide with R146^4.41^ and between the sulfonyl and K141^ICL2^. The indazole of VU6007678 forms a π-π interaction with F130^3.52^, a cation-π interaction with R134^3.56^ and at the bottom of TM3, and a hydrophobic interaction with Y129^3.51^. Other residues that make up the VU6007675 binding site include Y68^2.42^ and Y138^ICL2^. These extensive interactions likely explain why VU6007678 has high cooperativity with ACh and its high affinity for the M_5_ mAChR active state. Given the low affinity of VU6007678 for the inactive state M_5_ mAChR, as observed in radioligand binding with antagonist [^3^H]-NMS^29^, we hypothesized that these binding site residues undergo substantial rearrangement during activation. Superimposition of the ACh-VU6007678 structure with the inactive tiotropium-bound M_5_ mAChR crystal structure^18^ revealed significant conformational changes in the allosteric binding site. F130^3.52^, Y138^ICL2^, and K141^ICL2^ show marked inward movement toward the TM core, while R134^3.56^ moves outward (Fig. 4j). These coordinated conformational changes create the allosteric binding site (Supplementary Fig. 5e,f) enabling VU6007678 binding and explaining its state-dependent affinity profile.

We performed GaMD simulations on the ACh-VU6007678-bound M_5_ mAChR-mini-Gα_qiN_ complex to validate the binding pose of VU6007678 and examine its dynamic interactions with the receptor. Both ACh and VU6007678 maintained stable positions throughout the simulation, with ACh showing minimal fluctuations (RMSD = 1.70 ± 0.28 Å) and VU6007678 displaying moderate mobility while remaining in its binding pocket (RMSD = 4.85 ± 0.56 Å) (Fig. 4h, l).

The VU6007678 binding site contains several residues that vary across mAChR subtypes (Fig. 5a). While Y68^2.42^ is unique to M_5_, appearing as phenylalanine in other mAChRs, R134^3.56^ and R146^4.41^ are conserved among M_1_, M3, and M_5_ but differ in M_2_ and M_4_. V123^3.45^ varies between leucine, isoleucine, and valine across M_1_-M_4_, while I149^4.44^ alternates between leucine, methionine, and valine in these subtypes. The ICL2 lysine is conserved in M_1_-M_3_ but appears as arginine in M_4_. In contrast, F126^3.48^, F130^3.52^, T133^3.55^, Y138^ICL2^, and M150^4.45^ remain fully conserved across all five mAChR subtypes. This partially non-conserved binding site architecture provided an opportunity to further validate and characterize the binding site through mutagenesis studies and functional assessment using the TruPath G protein activation assay^46^.

**Figure 5:**
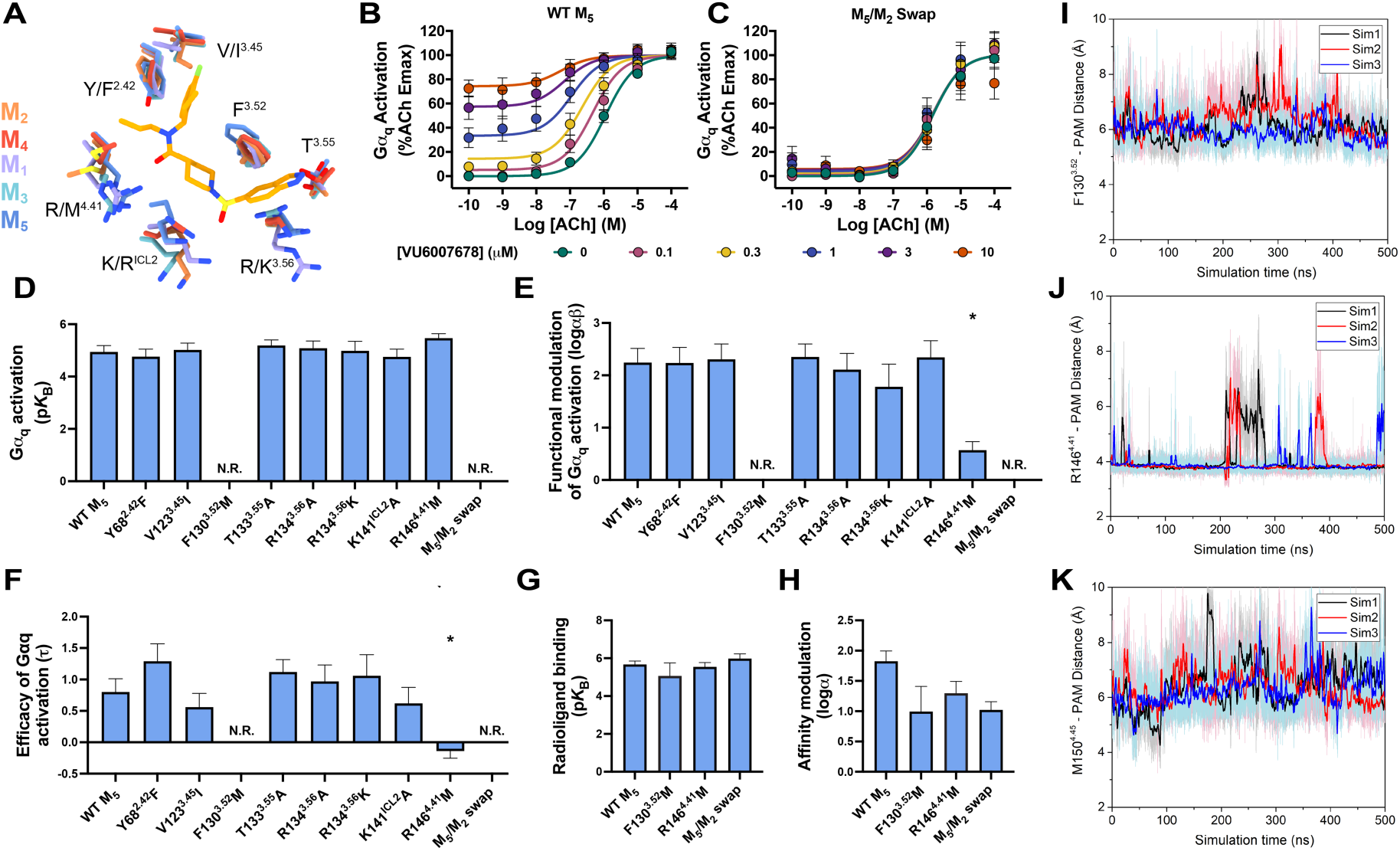
Structural and functional analysis of the VU6007678 allosteric binding site. **(A)** Comparison of VU6007678 allosteric binding site residues (stick representation) across the M_1_-M_5_ mAChRs. M_1_ mAChR (PDB: 6OIJ) is shown in purple, M_2_ mAChR (PDB: 6OIK) is shown in orange, M_3_ mAChR (PDB: 8E9Z) is shown in light blue, M_4_ mAChR (PDB: 7TRK) is shown in red and the M_5_ mAChR is shown in dark blue. **(B-C)** Interaction of ML380 with ACh in Trupath G protein activation assay in (B) WT M_5_ mAChR or (C) M_5_/M_2_ swap expressing CHO cells. Data points represent mean ± SEM of three to eight individual experiments performed in duplicate. WT M_5_ mAChR *n* = 8, M_5_/M_2_ swap mutant *n =* 3. An operational model of allosterism was fit to the data. **(D-F)** Key pharmacological parameters for the interaction of ACh and VU6007678 in Gα_q_ activation at WT M_5_ mAChR and mutants. Data points represent mean ± SEM of three to eight individual experiments performed in duplicate. WT M_5_ mAChR *n* = 8, M_5_/M_2_ swap, F130^3.52^M, T133^3.55^A, R134^3.56^A, R134^3.56^K mutants *n =* 3, Y68^2.42^F, V123^3.45^I, K141^ICL2^A, R146^4.41^M mutants *n* = 4. An operational model of allosterism was fit to the data. Parameters obtained are listed in Supplementary Table 4. **(G)** Affinity of VU6007678 (p*K*_B_) and **(H)** logarithm of affinity cooperativity (logα) between the orthosteric agonist (ACh) and allosteric modulator (VU6007876) at the WT M_5_ mAChR and selected mutants obtained in [^3^H]-NMS equilibrium radioligand binding studies between ACh, VU6007678 and [^3^H]-NMS. Data points represent mean ± SEM of three to five individual experiments performed in duplicate. WT M_5_ mAChR *n* = 3, F130^3.52^M, R146^4.41^M *n* = 4, M_5_/M_2_ swap *n* = 5. Data was fit to an allosteric ternary complex model. Parameters obtained listed in Supplementary Table 4. **(I-K)** Time courses of distances from VU6007678 to (I) F130^3.52^, (J) R146^4.41^, (K) M150^4.45^ in Å calculated from GaMD simulations performed with three separate replicates as indicated through different coloured traces.

In WT M_5_ mAChR, VU6007678 demonstrated robust positive functional cooperativity with ACh signalling and exhibited significant allosteric agonism (Fig. 5b) in a G_q_ TruPath experiment. Given the selectivity of VU6007678 for the M_5_ mAChR over the M_2_ mAChR and its extensive ICL2 interactions, we first investigated non-conserved residues at the bottom of TMs 3, 4, and ICL2 by mutating them to their M_2_ mAChR counterparts. A construct containing multiple mutations (S131^3.53^C, I132^3.54^V, R134^3.56^K, R139^ICL2^P, A140^ICL2^V, P144^4.39^T, R146^4.41^M, I149^4.44^M, G152^4.47^A, and L153^4.48^A (referred to as the M_5_/M_2_ swap) caused a complete loss in the ability of VU6007678 to modulate ACh signalling and to display allosteric agonism (Fig. 5c). Individual residue analysis revealed that the R146^4.41^M mutation reduced functional modulation and eliminated allosteric agonism, while the F130^3.52^M mutation completely abolished the affinity, functional modulation, and agonism of VU6007678 (Fig. 5d-f, Supplementary Fig. 6). These effects align with our structural data with R146^4.41^ forming a hydrogen bond with the carboxamide group of VU6007678, and a π-interaction between F130^3.52^ and the indanyl core of VU6007678 (Fig. 4i). Other single mutations (Y68^2.42^F, V123^3.45^I, T133^3.55^A, R134^3.56^A, R134^3.56^K, K141^ICL2^A, R146^4.41^M) showed no significant changes in affinity, functional modulation, or allosteric agonism (Fig. 5d-f, Supplementary Fig. 6).

Radioligand binding studies of F130^3.52^M, R146^4.41^M, and the M_5_/M_2_ swap showed no significant change in VU6007678 affinity (Fig. 5g, Supplementary Fig. 7, Table 4). While the binding cooperativity (log α) was reduced in these constructs (Fig. 5h, Supplementary Fig. 7, Table 4), the values were not significantly different from WT M_5_ mAChR, likely due to higher uncertainty in the parameter calculations for the F130^3.52^M and M_5_/M_2_ swap constructs. The degree of efficacy modulation (β) by VU6007678 on ACh can be calculated by subtracting the binding modulation (log α) from the functional modulation (log αβ)^34^. This calculation at WT M_5_ mAChR yielded a log β value of 0.4 ± 0.2, indicating that VU6007678’s allosteric effect is primarily mediated through binding cooperativity, with a smaller contribution from efficacy modulation. This value could only be compared to the R146^4.41^M mutant, as the F130^3.52^M and M_5_/M_2_ swap showed no functional modulation. The R146^4.41^M mutant yielded a log β value of −0.7 ± 0.3, suggesting impaired efficacy modulation by VU6007678. These findings are supported by our previous characterisation of VU6007678 in receptor alkylation studies, which revealed modest efficacy modulation in functional IP one assays^29^. Interestingly, this observation was unique to VU6007678 in the structure-activity relationship (SAR) study. Collectively, these data highlight the importance of the VU6007678 binding site residues in mediating both functional and efficacy modulation, possibly due to the binding site’s location overlapping the DRY activation motif and its proximity to the G protein binding site.

Further validation of the allosteric binding site through GaMD simulations revealed key interactions between VU6007678 and receptor residues. The PAM formed stable interactions with F130^3.52^, R146^4.41^, and M150^4.45^ (Fig. 5i-k), consistent with both the cryo-EM structure and mutagenesis data. In contrast, hydrogen bond interactions between VU6007678 and receptor residues R134^3.56^, T133^3.55^, and K141^ICL2^ showed greater variability with larger distances and higher fluctuations (Supplementary Fig. 8), aligning with our experimental observations where mutations of these residues did not significantly affect VU6007678 function. These results highlight how VU6007678 engages with residues involved in the inactive-to-active transition of the M_5_ mAChR, particularly through stable interactions with F130^3.52^ and R146^4.41^, while maintaining more dynamic interactions with K141^ICL2^. Together, this suggests VU6007678 helps stabilize key residues involved in receptor activation.

## Discussion

Allosteric modulators for the mAChRs have long been pursued for selective targeting of a specific mAChR subtype. Whilst selective allosteric modulators for the M_5_ mAChR have been discovered, the development and application of these have lagged compared to allosteric modulators selective for other mAChRs, particularly the M_1_ and M_4_ mAChR. This partly reflects the limited structural knowledge of the M_5_ mAChR overall, as well as the specific lack of insight into the allosteric binding site for M_5-_selective PAMs. Here we present a cryo-EM structure of the M_5_ mAChR bound to the orthosteric agonist iperoxo that ‘completes’ the active state structure ensemble for all five mAChR subtypes. Initial attempts to solve an allosteric modulator co-bound structure with ML380 and iperoxo were unsuccessful. Yet through use of analytical pharmacology to determine the optimum orthosteric and allosteric ligand combination and improved biochemical techniques, we obtained a high-resolution structure of the M_5_ mAChR co-bound to the endogenous orthosteric agonist ACh and the selective PAM VU6007678. Our data begins to explain several facets of selective allosteric modulation at the M_5_ mAChR including 1) the historic difficulties in developing subtype selective PAMs for this mAChR subtype, 2) how changes to ML380 led to the improved PAM VU6007678, 3) subtype selectivity, and 4) mechanism of action.

The *in vivo* translation of M_5_ mAChR selective PAMs has been plagued by issues relating to DMPK and suboptimal partition coefficients (K_p_)^25^. The observation that the allosteric binding site for VU6007678 is in the TM bundle partially explains this, as to reach this allosteric binding site, modulators must display a high degree of lipophilicity. Despite this, the VU6007678 scaffold offers numerous opportunities for modifications, and our structure will aid the process of developing improved allosteric modulators as it provides information on the molecular interactions that occur between VU6007678 and the M_5_ mAChR at its allosteric binding site. Specifically, Y68^2.42^ is the only residue within the allosteric binding site that is different at all M_1_-M_4_ mAChR subtypes. It may therefore be possible to introduce and optimize the presence of various polar functional groups on the indanyl core to promote hydrogen bonding between the ligand and receptor and enhance affinity whilst reducing the compound’s lipophilicity. Note, the compounds reported in the previous SAR series all had a fluorine functional group attached to the indanyl core^26^. Enhancing the selectivity of PAMs for the M_5_ mAChR will be, in part, crucial to the future clinical success of M_5_ mAChR PAMs. All M_5_ PAMs discovered to date display activity at the M_1_ and M_3_ mAChRs whilst they are most selective against the M_2_ and M_4_ mAChR where they display very little to no activity. Analysis of the residues within the VU6007678 allosteric binding site may explain the basis for this as R134^3.56^ and R146^4.41^ are fully conserved between the M_5_ mAChR and M_1_/M_3_ mAChRs and non-conserved between the M_5_ mAChR and the M_2_/M_4_ mAChRs.

Despite the absence of an ML380-bound M_5_ mAChR structure, given that VU6007678 is a derivative of ML380, predictions on how ML380 interacts with this site can be made. This allows for an explanation of how the changes to VU6007678 led to an improved PAM. The extension of the ethyl present in ML380 into a propyl group at VU6007678 gives rise to an extra interaction with I149^4.44^. In addition, the substitution of the trifluoromethylbenzyl group at ML380 with the indanyl group gives rise to more hydrophobic interactions and greater engagement with the top of the allosteric site consisting of V123^3.45^, F126^3.48^ and M150^4.45^ where edge-to-face π-interaction takes place with F126^3.48^.

Our structure offers a few observations on the mechanism of action for PAMs at the M_5_ mAChR. First, the allosteric binding site of VU6007678 is in close proximity to the highly conserved DRY activation motif, indicating that PAM binding at this allosteric site stabilizes the active state of the M_5_ mAChR. Second, the mAChR family has served as a model receptor family for the study of allosteric modulation at GPCRs^19^. Despite a wealth of mAChR allosteric modulators with different scaffolds available, this is only the second site to be confirmed by structural biology studies at the mAChRs^33,34^ (Fig. 6a). All PAMs discovered to date bind to the mAChRs at what is termed the common ECV allosteric site in a solvent accessible vestibule on top of the orthosteric binding site. Here PAMs exert their mechanism of action through two mechanisms, trapping the orthosteric ligand in the orthosteric site through stabilising the active state and sterically hindering the orthosteric ligand dissociation^47,48^. Due to the distal location of this allosteric binding site to the mAChR orthosteric site, this would indicate PAMs exert their mechanism of action through this site only by stabilising the active state. Such a mechanism would be akin to that observed for other allosteric modulators binding to the same site at different GPCRs^49^, including but not limited to, compound 6FA at the β_2_AR^50^ and LY3154207 at the D1R^51–53^. The discovery of a methionine (M^4.45^) within the VU6007678 allosteric binding site allows for potential methionine labelling of this residue. Subsequent characterisation of PAM activity at a methionine labelled M_5_ mAChR through spectroscopic approaches will facilitate further exploration of how PAMs exert their mechanism of action at the M_5_ mAChR as was done for LY2119620 at the M_2_ mAChR^54^.

**Figure 6:**
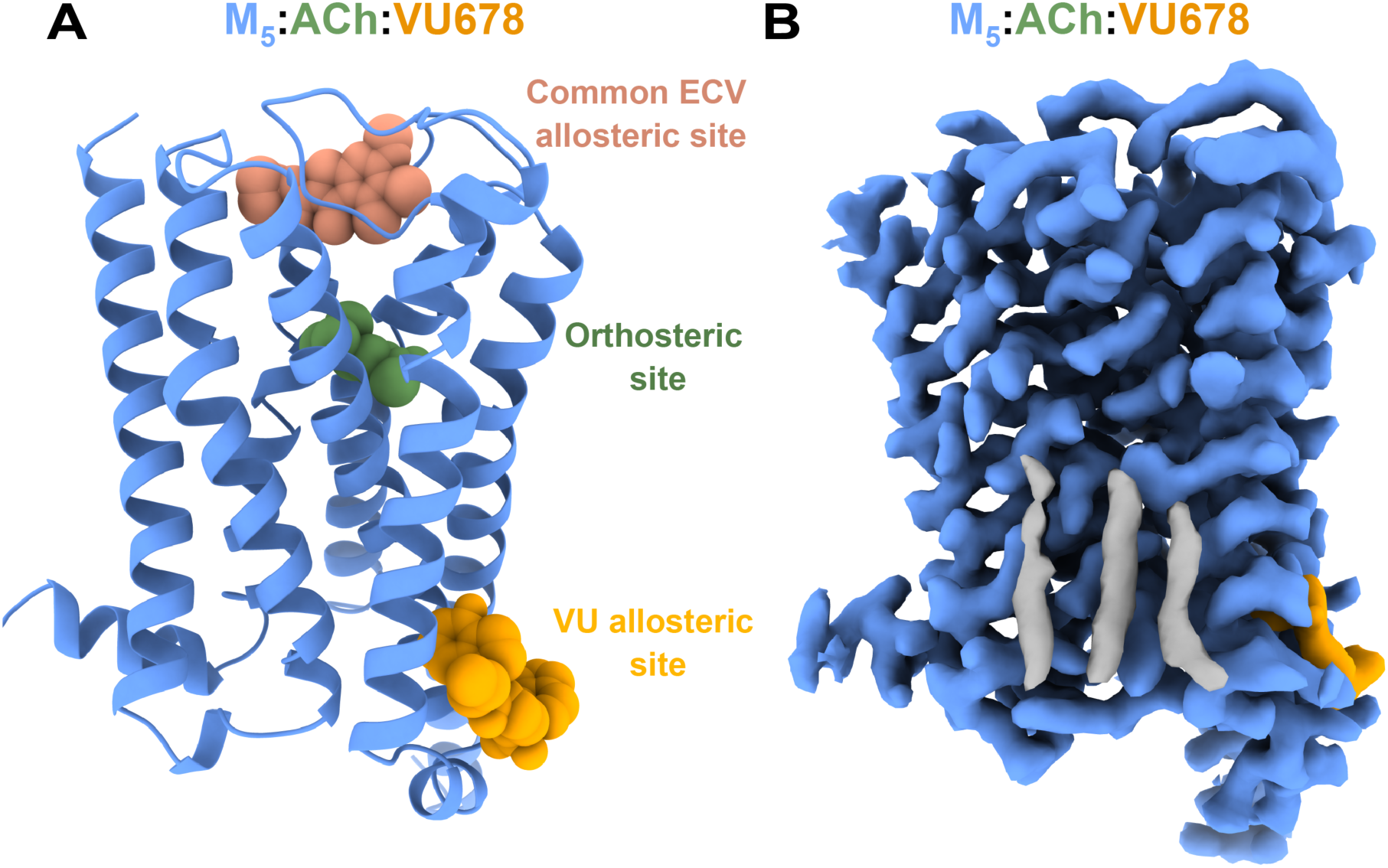
Allosteric sites at the mAChRs. **(A)** Model of the ACh-VU6007678 bound M_5_ mAChR showing the novel allosteric binding site discovered for VU6007678 (orange spheres) and the ‘common’ ECV allosteric site occupied by LY2119620 (salmon spheres). ACh is shown as green spheres in the orthosteric binding site. **(B)** Cryo-EM density (contour level 0.47) for lipid molecules observed at the bottom of TM2,3,4 in the orthosteric binding site.

At the bottom of the TM2,3,4 interface of the ACh-VU6007678 structure we observed well defined densities that likely correspond to three cholesterol molecules (Fig. 6b). Note in our iperoxo structure this site was occupied by partial density that we cautiously hypothesised could be ML380. Considering the much higher resolution of the ACh-VU6007678 structure, and assignment of the VU6007678 binding site at the bottom of the extrahelical interface at TMs3-4, it is likely that the density present in the iperoxo-bound structure represents cholesterol molecules. Especially given that this site is recognised as a common cholesterol site within class A GPCRs^44,55^ and that neurosteroids and steroid hormones, including derivatives of cholesterol, have displayed modulatory properties at the M_5_ mAChR^56,57^. Due to the lower density and map quality in this region, conclusive assignment of this density to either ML380 or cholesterol was difficult in the iperoxo structure, highlighting the map quality that is required to conclusively assign density to small molecules. Particularly at extrahelical regions where a number of lipid and cholesterol molecules may be present.

Altogether and more broadly, the discovery of this novel allosteric binding site at the M_5_ mAChR adds to the knowledge of allostery at the mAChR, specifically on how allosteric modulators engage with the mAChRs, and provides an additional avenue through which to target these highly conserved proteins.

## Materials and Methods

### Receptor & G protein co-expression

A modified M_5_ mAChR construct was used where residues 237-421 of ICL3 were removed and HA signal sequence and anti-Flag epitope tag were added to the N terminus. Modified M_5_ mAChR was fused to a mini-Gα_qiN_ chimeric construct that is a mini-Gα_s_ substituted with Gα_q_ residues at the receptor interface and the αN of Gα_i_^38,39^. A 3C protease recognition site and GGGS linker was used to separate the receptor and G protein. The fused M_5ΔICL3_mGα_sQi_/construct was cloned into a pFastbac baculovirus transfer vector. G protein β_1_ and γ_2_ subunits were cloned into a pVL1392 baculovirus transfer vector with the β subunit modified to contain a carboxy (C)-terminal 8× histidine tag. *Trichoplusia ni* (Hi5) insect cells were grown in ESF 921 serum-free media (Expression Systems) and infected at a density of 4.0 × 10^6^ cells per millilitre with a 1:1 ratio of M_5ΔICL3_mGα_sQi_ to Gβ_1_γ_2_ viruses and shaken at 27°C for 48-60 hours. Cells were harvested by centrifugation and cell pellets were flash frozen using liquid nitrogen and stored at −80°C.

### Single chain stabilising fragment expression and purification

A single-chain construct of Fab16 (scFv16), tagged with an 8× histidine sequence at the C-terminus, was cloned into a modified pVL1392 baculovirus transfer vector for secreted expression in *Trichoplusia ni* (Hi5) insect cells (Expression Systems). The cells were cultured in serum-free ESF 921 media (Expression Systems), infected at a density of 4.0 × 10^6^ cells per millilitre, and incubated with shaking at 27°C for 48-72 hours. For purification, the pH of the supernatant from the baculovirus-infected cells was adjusted with Tris pH 8.0. Chelators were neutralized by adding 5 mM CaCl_2_ and stirring the solution for 1 hour at 25°C. Precipitates were cleared by centrifugation, and the supernatant was then applied to Ni-NTA resin. The column was washed with a solution of 20 mM Hepes pH 7.5, 50 mM NaCl, and 10 mM imidazole, followed by a second wash containing the same buffer with 100 mM NaCl. The scFv16 protein was eluted using the low-salt buffer with 250 mM imidazole. SDS-PAGE with Coomassie staining was used to assess purity of the eluted protein. Finally, the sample was concentrated, flash-frozen in liquid nitrogen, and stored at −80°C.

### Nb35 expression and purification

Nb35 was expressed in the periplasm of the BL21(DE3) Rosetta *Escherichia coli* cell line using an autoinduction approach^58^. Transformed cells were cultured at 37°C in a modified ZY medium containing 50 mM phosphate buffer (pH 7.2), 2% tryptone, 0.5% yeast extract, 0.5% NaCl, 0.6% glycerol, 0.05% glucose, and 0.2% lactose, with 100 μg/ml carbenicillin and 35 μg/ml chloramphenicol. When the culture reached an OD600 of 0.7, the temperature was reduced to 20°C for approximately 16 hours, after which cells were harvested by centrifugation and stored at −80°C. To purify, cells were lysed in ice-cold buffer (0.2 M Tris pH 8.0, 0.5 M sucrose, 0.5 M EDTA) at a ratio of 1g of cell pellet to 5mL of lysis buffer for 1 hour at 4°C. 2x volume of ice cold MQ was added and incubated for an additional 45 minutes at 4°C. Lysate was centrifuged to remove cell debris and supernatant containing Nb35 was spiked with 20 mM HEPES pH 7.5, 150 mM NaCl, 5 mM MgCl_2_, 5 mM Imidazole and applied to Ni-NTA resin followed by 90 minutes incubation at 4°C. The column was washed with 40 column volumes (CV) of wash buffer (20 mM HEPES pH 7.5, 500 mM NaCl, 5 mM Imidazole) and eluted with elution buffer (20 mM HEPES pH 7.5, 100 mM NaCl, 250 mM Imidazole). Elute was concentrated and stored at 80°C.

### Complex purification

#### Iperoxo-bound M_5_ mAChR-mini-Gα_qiN_ complex

M_5ΔICL3_ mGα_sQi_ co-expressed with Gβ_1_γ_2_ was thawed and lysed in in 20 mM HEPES pH 7.4, 5 mM MgCl_2_, 1 μM Ipx and protease inhibitors (500 µM PMSF, 1 mM LT, 1 mM benzamidine). The sample was rotated at room temperature for 15 minutes and spiked with apyrase towards the end of the 15 minutes. The pellet was spun down by centrifugation and solubilized in 20 mM HEPES pH 7.4, 100 mM NaCl, 5 mM MgCl_2_, 5 mM CaCl_2_, 0.5% LMNG (Anatrace, Maumee, OH, USA), 10 μM Ipx, and protease inhibitors (500 µM PMSF, 1 mM LT, 1 mM benzamidine). The resuspended pellet was homogenised in a Dounce homogeniser and complex formation was initiated through addition of scFv16. The sample was incubated with stirring at 4°C for 2 hours followed by centrifugation to remove insoluble material. Solubilised complex was bound to equilibrated M1 anti-Flag affinity resin though batch binding at room temperature for 1 hour. The resin was packed into a glass column and washed with 20 mM HEPES pH 7.4, 100 mM NaCl, 5 mM MgCl_2_, 5 mM CaCl_2_, 0.01% LMNG, and 1 μM Ipx until no more protein was coming off the column as determined by Bradford. Complex was eluted using 20 mM HEPES pH 7.4, 100 mM NaCl, 5 mM MgCl_2_, 0.01% LMNG, and 1 μM Ipx in the presence of 5 mM EDTA and 0.1 mg/ml Flag peptide. Eluted complex was concentrated in an Amicon Ultra-15 100 kDa molecular mass cut-off centrifugal filter unit (Millipore, Burlington, MA, USA) and purified by size exclusion chromatography (SEC) on a Superdex 200 Increase 10/300 GL (Cytiva, Marlborough, MA, USA) in 20 mM HEPES pH 7.4, 100 mM NaCl, 5 mM MgCl_2,_ 0.01% LMNG, and 1 μM Ipx. Fractions containing complex (as determined by SDS-page and Coomassie staining) were pooled and concentrated to 3.5 mg/mL, flash frozen using liquid nitrogen and stored at −80°C.

#### ACh-VU6007678-bound M_5_ mAChR-mini-Gα_qiN_ complex

The ACh-VU6007678 M_5_ mAChR sample was purified in an identical manner with the following changes; 1) complex formation was initiated through addition of scFv16 and Nb35, 2) ACh was included in all buffers at 100 μM until SEC where a concentration of 10 μM was utilised, 3) VU6007678 was present throughout the purification at a concentration of 10 μM. The final purified product was concentrated to 18 mg/mL.

### Vitrified sample preparation and data collection

For the iperoxo-M_5_ mAChR sample, EMAsian - TiNi 200 mesh 1.2/1.3TiNi 200 mesh 1.2/1.3 grids were glow discharged using Pelco EasyGlow for 90 seconds with 15-mA current. Prior to grid freezing, the iperoxo-M_5_ mAChR sample was spiked with 30 μM of ML380 and incubated ON at 4°C. A 3 μL of this spiked sample was applied and flash frozen in liquid ethane using a Vitrobot markIV with a blot force of 4 and blot time of 2 seconds at 100 % humidity and 4 °C. Data were collected on a Titan Krios (Thermo Fisher Scientific) 300 kV electron microscope equipped with a K3 detector, 50 μm C2 aperture, no objective aperture inserted, indicated magnification x 130 000 in nanoprobe TEM mode, a slit width of 10eV, pixel size 0.65 Å, exposure rate 10.57 counts per pixel per second, exposure time 2.68 s, total exposure 60 e Å^-2^, and 60 frames. In total, 9104 movies were collected.

For the ACh-VU6007678 M_5_ mAChR sample, 3 µL of sample was applied to glow-discharged (15 mA, 180 s) UltrAufoil R1.2/1.3 300 mesh holey grid (Quantifoil) and were frozen in liquid ethane using a Vitrobot mark IV (Thermo Fisher Scientific) at 100% humidity and 4°C with a blot time of 2 s and blot force of 10. The sample was collected similarly, except at 105kX magnification with a pixel size of 0.82 Å, and 7489 movies were collected.

### Image Processing

For the iperoxo-M_5_ mAChR sample, 9104 movies were collected and adjusted for beam-induced motion by MotionCor2^59^. Non-dose weighted micrographs were used for CTF estimation using Gctf^60^. 8341 micrographs were identified as having a ctf fit resolution below 4 Å, and these were selected for further examination. 3,833,583 particles were autopicked using Gautomatch (https://www2.mrc-lmb.cam.ac.uk/research/locally-developed-software/zhang-software/#gauto). The particles were extracted with relion-3.1^61^ and then imported into CryoSparc^62^ for rounds of 2D classification, ab initio 3D and 3D refinement to obtain a 3.04 Å model. Particles were taken to Relion3.1 for polishing and subsequent 3D refinement back in Cryosparc yielded a final model of 2.75 Å based on the gold standard Fourier shell correlation cut-off of 0.143 from 426,714 particles. A further local refinement was performed to generate a receptor-focused map (2.67 Å).

The ACh-VU6007678 M_5_ mAChR sample was processed similarly. 7489 movies were collected and adjusted for beam-induced motion by MotionCor2^59^. Non-dose weighted micrographs were used for CTF estimation using Gctf^60^. 4,436,835 particles were autopicked using Gautomatch (https://www2.mrc-lmb.cam.ac.uk/research/locally-developed-software/zhang-software/#gauto). The particles were extracted with relion-3.1^61^ and then imported into CryoSparc^62^ for rounds of 2D-classification and heterogenous refinement. A set of particles (∼900k) were polished in relion3.1 and a final round of 3D-classification (no alignment) was performed. This final set of 418,794 particles were finally subjected to non-uniform refinement with CTF-refinement in CryoSparc, resulting in a final map of 2.06 Å based on the gold standard Fourier shell correlation cut-off of 0.143. A further local refinement was performed to generate a receptor-focused map (2.11 Å).

### Model building and refinement

An initial receptor model was generated from the cryo-EM structure of the M_4_ mAChR receptor (PDB: 7TRP). An initial model for the G protein (mini-Gα_qiN_:Gβ_1_Gγ_2_:ScFv16) was generated from the CCK1:mini-Gα_qiN_ complex (PDB: 7MBY). Initial models were placed in the EM maps using UCSF ChimeraX^63^ and rigid-body-fit using PHENIX^64^. Models were refined with iterative rounds of manual model building in Coot^65^ and ISOLDE, and real-space refinement in PHENIX. Ligands ACh and iperoxo were obtained from the monomer library, while initial model and restraints for VU6007678 were generated using the GRADE web server (https://grade.globalphasing.org). Model validation was performed with MolProbity^66^ and the wwPDB validation server^67^. Figures were generated with UCSF ChimeraX and PyMOL (Schrödinger).

### Cell culture

FlpIn Chinese hamster ovary (CHO) cells (Thermo Fisher Scientific) stably expressing M_5_ mAChR constructs were cultured at 37°C in 5% CO_2_ using Dulbecco’s modified Eagle’s medium (DMEM; Invitrogen) supplemented with 5% foetal bovine serum (FBS; ThermoTrace). At confluence, media was removed, and cells were washed with phosphate-buffered saline (PBS) and harvested from tissue culture flasks using versene (PBS with 0.02% EDTA). The cells were pelleted by centrifugation at 350g for three minutes and then resuspended in DMEM with 5% FBS. Subsequently, the cells were either plated for an assay or reseeded into a tissue culture flask.

### Inositol Monophosphate (IP1) Accumulation Assay

FlpIn CHO cells stably expressing either WT or mutant hM_5_ mAChR were seeded in clear, flat-bottom 96-well plates at a density of 10,000−25,000 cells per well (depending on the cell line) one day prior to the assay. The optimal cell density for each line was chosen based on achieving an IP1 response that fell within the linear range of the IP1 standard curve. On the assay day, the medium was replaced with stimulation buffer (Hanks’ balanced salt solution (HBSS) containing 10 mM HEPES, 1.3 mM CaCl_2_, and 30 mM LiCl, pH 7.4) and allowed to incubate for 60 minutes at 37°C before ligand stimulation. After this pre-incubation, the buffer was replaced, and cells were exposed to ligands for 60 minutes at 37°C in a 5% CO_2_ atmosphere, with a total assay volume of 100 μL. Following the 60-minute stimulation, ligands were removed by rapid removal of buffer. Cells were lysed by freeze-thawing in 30 μL of stimulation buffer. IP1 accumulation was then quantified using the HTRF IP-One assay kit (Cisbio), with fluorescence measured on an EnVision multilabel plate reader (PerkinElmer).

### TruPath – G protein Activation Assay

Upon reaching 60-80% confluence, FlpIn CHO cells stably expressing WT or mutant hM_5_ mAChR were transiently transfected using Polyethylenimine (PEI; Sigma-Aldrich). For each well, 10 ng of each plasmid (pcDNA5/FRT/TO-Gα_i1_-RLuc8, pcDNA3.1-β_3_, and pcDNA3.1-Gγ_9_- GFP2) was added in a 1:1:1 ratio, totalling 30 ng. These plasmids were generously provided by Prof. Bryan Roth from the University of North Carolina. The cells were then plated at 30,000 cells per well into 96-well Greiner CELLSTAR white-walled plates (Sigma-Aldrich). After 48 hours, the cells were washed with 200 μL PBS and replaced with 1x HBSS supplemented with 10 mM HEPES. The cells were incubated for 30 minutes at 37°C before adding 10 μL of 1.3 μM Prolume Purple coelenterazine (Nanolight Technology, Pinetop, AZ). Following a further 10-minute incubation at 37°C, bioluminescence resonance energy transfer (BRET) measurements were performed using a PHERAstar FSX plate reader (BMG Labtech) with 410/80-nm and 515/30-nm filters. Four baseline measurements were taken before adding drugs or vehicle, bringing the final assay volume to 100 μL, followed 10 more minutes of readings. The BRET signal was calculated as the ratio of 515/30-nm emission to 410/80-nm emission. This ratio was vehicle-corrected using the initial four baseline measurements and then baseline-corrected again using the vehicle-treated wells. Data were normalized to the maximum ACh response to allow for grouping of results.

### Radioligand Binding

FlpIn CHO cells stably expressing WT hM_5_ mAChR or mutants were plated at 25,000 cells per well in 96-well isoplates (PerkinElmer Life Sciences) and incubated overnight at 37°C in a 5% CO_2_ incubator. The following day, the cells were washed with PBS and incubated in 20 mM HEPES, 100 mM NaCl, 10 mM MgCl_2_, pH 7.4. For saturation binding experiments, the cells were incubated with varying concentrations of the orthosteric antagonist [^3^H]-*N*- methylscopolamine ([^3^H]-NMS; specific activity, 70 Ci/mmol, Perkin Elmer) in a final volume of 100 μL for 6 hours at room temperature. For interaction experiments between orthosteric agonist and allosteric modulator, competition binding was performed between a K_D_ concentration of [^3^H]-NMS and varying concentrations of an orthosteric drug in the presence of different concentrations of an allosteric modulator, in a total volume of 100 μL binding buffer. For all experiments, non-specific binding was defined using 10 μM of atropine. The assay was terminated by the rapid removal of the radioligand, followed by two 100 μL washes with ice-cold 0.9% NaCl buffer. Radioactivity was measured by adding 100 μL of Optiphase Supermix scintillation fluid (Perkin Elmer) and counted using a MicroBeta^2^ Plate Counter (PerkinElmer Life Sciences).

### Data Analysis

All data were analyzed using GraphPad Prism 10 (GraphPad Software, San Diego, CA). The interaction between orthosteric agonist and allosteric modulator in functional assays was analysed using an operational model of allosterism to determine functional modulation (log αβ) and affinity (p*K*_B_) parameters^35^. Radioligand saturation binding experiments with [^3^H]-NMS to determine B_max_ and p*K*_D_ values were determined as previously described^31^. For the radioligand binding interaction of orthosteric agonist with various concentrations of allosteric modulator, the data were fit to an allosteric ternary complex model to derive p*K*_B_ and α binding cooperativity parameters^68^. All affinity, potency, cooperativity, and efficacy parameters were estimated as logarithms. Statistical analysis between different treatment conditions was performed using one-way ANOVA, with a p-value of < 0.05 considered significant.

### Gaussian accelerated Molecular Dynamics (GaMD)

GaMD is an enhanced sampling method that works by adding a harmonic boost potential to reduce the system energy barriers^69,70^. When the system potential 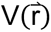 is lower than a reference energy E, the modified potential 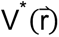 of the system is calculated as:

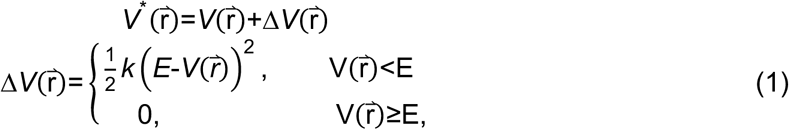

where *k* is the harmonic force constant. The two adjustable parameters E and k are automatically determined on three enhanced sampling principles. First, for any two arbitrary potential values 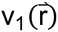 and 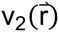 found on the original energy surface, if 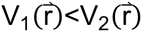, Δ*V* should be a monotonic function that does not change the relative order of the biased potential values; i.e., 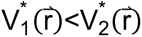. Second, if 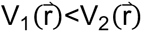, the potential difference observed on the smoothened energy surface should be smaller than that of the original; i.e., 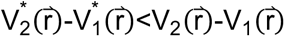. By combining the first two criteria and plugging in the formula of 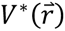 and Δ*V*, we obtain:

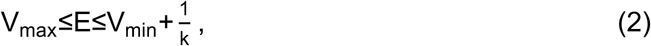

Where V_min_ and V_max_ are the system minimum and maximum potential energies. To ensure that Eq. 2 is valid, *k* has to satisfy: k≤1/(V_max_-V_min_). Let us define: k=k_0_·1/(V_max_-V_min_), then 0<k_0_≤1. Third, the standard deviation (SD) of Δ*V* needs to be small enough (i.e. narrow distribution) to ensure accurate reweighting using cumulant expansion to the second order: σ_ΔV_=k+E-V_avg_,σ_V_≤σ_0_, where V_avg_ and σ_V_ are the average and SD of ΔVwith σ_0_ as a user-specified upper limit (e.g., 10k_B_T) for accurate reweighting. When E is set to the lower bound E=V_max_ according to Eq. 2, k_0_ can be calculated as:

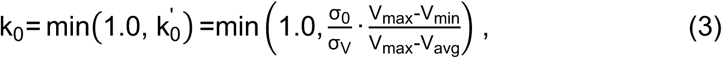

Alternatively, when the threshold energy E is set to its upper bound E=V_min_+1/k, k_0_ is set to:

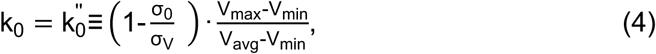

If 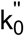 is calculated between 0 and 1. Otherwise, k_0_ is calculated using Eq. 3.

### GaMD Simulations and Simulation Analysis

The cryo-EM structures of the ACh-VU6007678-bound M_5_ mAChR-mini-Gα_qiN_ complex and the Iperoxo-bound M_5_ mAChR-mini-Gα_qiN_ complex were used to set up simulation systems. The initial model of the Iperoxo-ML380-bound M_5_ mAChR-mini-Gα_qiN_ complex was created by docking ML380 into the bottom of the TM2,3,4 interface. As in our previous studies^34^, the intracellular loop 3 (ICL3) of the receptor and the α-helical domain of the G protein, which were missing in the cryo-EM structure, were not modeled. The MD simulation systems were prepared by inserting the ACh-VU6007678-bound M_5_ mAChR-mini-Gα_qiN_ and the Iperoxo-bound M_5_ mAChR-mini-Gα_qiN_ complexes into a POPC (palmitoyl-2-oleoyl-sn-glycero-3- phosphocholine) lipid bilayer using VMD (Visual Molecular Dynamics). In each simulation system, the protein and lipid bilayer were solvated with TIP3P water molecules in a box of 12.5 nm x 12.5 nm x 14.0 nm with the periodic boundary condition. The system charge was neutralized with 150 mM NaCl. The AMBER FF14SB force field^71^ was applied for the proteins, while the AMBER LIPID21 force field^72^ was used for the lipids. The general amber force field (GAFF2) parameters^73^ for ACh, Iperoxo, ML380, and VU6007678 were generated with ANTECHAMBER. The two simulation systems were first energy minimized for 5,000 steps with constraints on the heavy atoms of the proteins and phosphor atom of the lipids. The hydrogen-heavy atom bonds were constrained using the SHAKE algorithm and the simulation time step was set to 2.0 fs. The particle mesh Ewald (PME) method^74^ was employed to compute the long-range electrostatic interactions and a cutoff value of 9.0 Å was applied to treat the non-bonded atomic interactions. The temperature was controlled using the Langevin thermostat with a collision frequency of 1.0 ps^-1^. Each system was equilibrated using the constant number, volume, and temperature (NVT) ensemble at 310K for 250 ps and under the constant number, pressure, and temperature (NPT) ensemble at 310 K and 1 bar for another 1 ns with constraints on the heavy atoms of the protein, followed by 10 ns short conventional MD (cMD) without any constraint.

The GaMD module implemented in the GPU version of AMBER18^69,70,75^ was then applied to simulate the ACh-VU6007678-bound M_5_ mAChR-mini-Gα_qiN_ and Iperoxo-bound M_5_ mAChR- mini-Gα_qiN_ complexes. The GaMD simulations included an 8-ns short cMD run to collect the potential statistics for calculating GaMD acceleration parameters, followed by a 56-ns GaMD equilibration after adding the boost potential. Finally, three independent 500-ns GaMD production simulations were conducted for each system with randomized initial atomic velocities. The average and standard deviation (SD) of the system potential energies were calculated every 800,000 steps (1.6 ns). All GaMD simulations were performed at the “dual-boost” level, where the reference energy was set to the lower bound. One boost potential was applied to the dihedral energetic term and the other to the total potential energetic term. The upper limit of the boost potential SD (σ_0_) was set to 6.0 kcal/mol for both the dihedral and the total potential energetic terms.

For each system, the three GaMD production trajectories were combined for analysis. The CPPTRAJ^76^ was applied to calculate the time-courses of the root-mean-square derivations (RMSDs) of agonists and PAMs relative to the simulation starting structure, as well as the distances between VU6007678 and key interacting residues in the receptor, including F130^3.52^, R146^4.41^, M150^4.45^, F126^3.48^, R134^3.56^, V123^3.45^, T133^3.55^ and K141^ICL2^.

## Acknowledgements

This work was funded by the Australian Research council (ARC) discovery Project DP190102950 (CV, AC) and a National Health and Medical Research Council of Australia (NHMRC) Program Grant APP1150083 (AC), NHMRC Project Grant APP1138448 (DMT), an NHMRC early career investigator Grant APP1196951 (DMT), a Discovery Early Career Researcher Award DE170100152 (DMT), and a US National Institutes of Health Grant R01GM132572 (YM). PRG was a Sir Keith Murdoch Fellow of the American Australian Association. CWL was funded by the William K. Warren Foundation. Cryo-EM imaging and sample vitrification were carried out at the Monash University Ramaciotti Centre for Cryo-Electron Microscopy. Data processing and storage of the cryo-EM datasets were supported by the Monash University MASSIVE high-performance computing facility and its supercomputing resources. This work used supercomputing resources with allocation awards TG-MCB180049 and BIO210039 through the US NSF-funded Advanced Cyberinfrastructure Coordination Ecosystem: Services & Support (ACCESS) program and project M2874 through the US DOE National Energy Research Scientific Computing Center (NERSC).

## Author Contributions

DMT designed the overall research. WACB, JIM, DMT designed, expressed, and purified protein samples. JIM, HV performed sample vitrification and cryo-EM imaging. WACB, JIM, DMT processed the EM data and generated and analysed atomic models. JW, KJ, YM designed, performed, and analysed MD simulations. WACB, BR, PRG, MY generated DNA constructs and performed pharmacology experiments. WACB, BR, PRG, MY, AC, CV, and DMT analysed pharmacology data. AMB and CWL provided chemical tools. YM, AC, CV, and DMT provided supervision. WACB, JIM, DMT wrote the manuscript with contributions and input from all authors.

