## Supplemental Data for "Cryo-EM reveals a new allosteric binding site at the M_5_ mAChR"

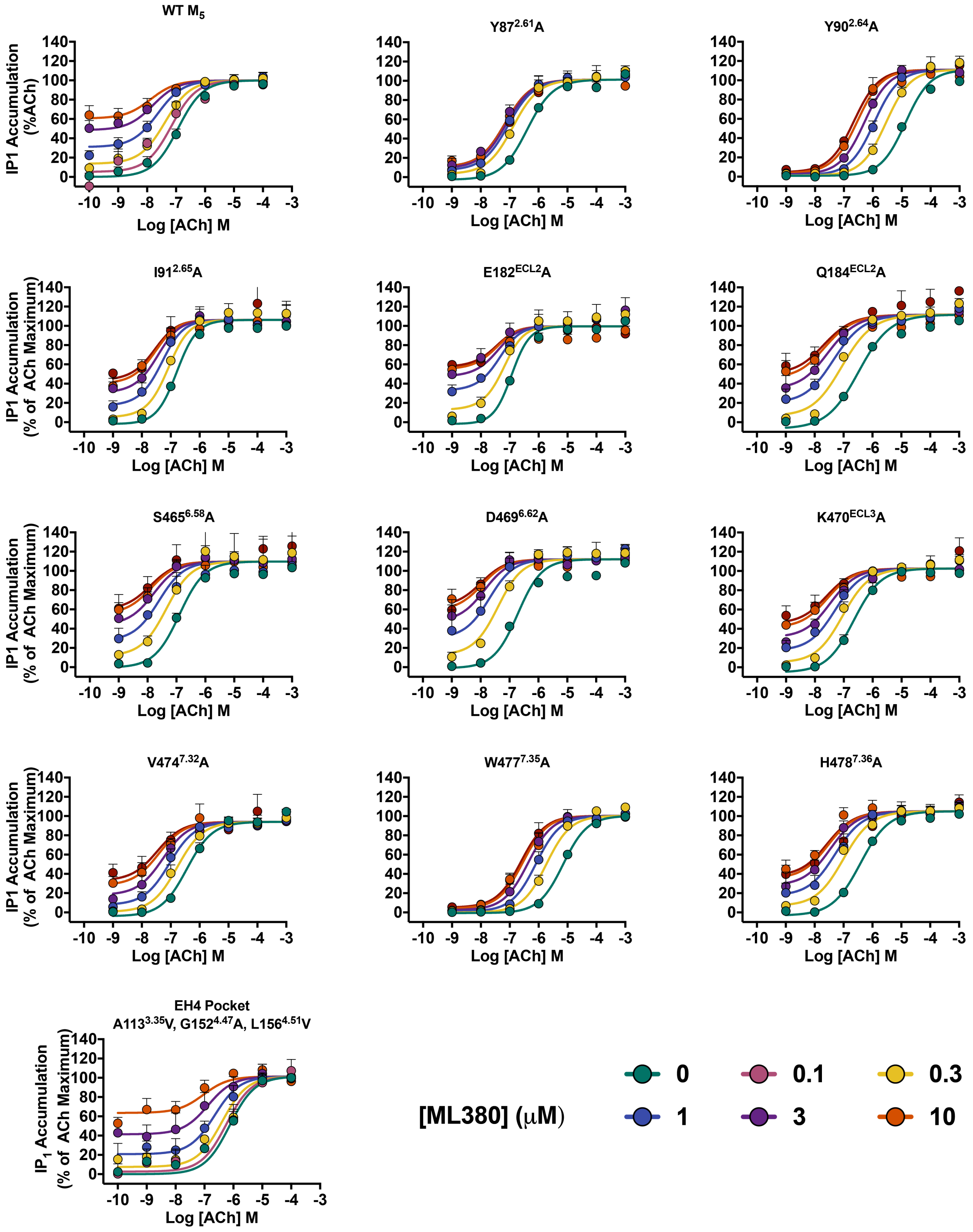


**Supplementary Figure 1. Interaction of ML380 with ACh at 'common' ECV allosteric site mutants.** Interaction of ML380 with ACh in an IP1 accumulation assay at 'common' ECV allosteric site mutant M$_5$ mAChRs expressed in CHO cells. Data points represent mean ± SEM of three individual experiments performed in duplicate. An operational model of allosterism was fit to the data. Parameters obtained are listed in Supplementary Table 1. Graphs for WT M$_5$ mAChR and EH4 pocket were reproduced from main Figure 1A.

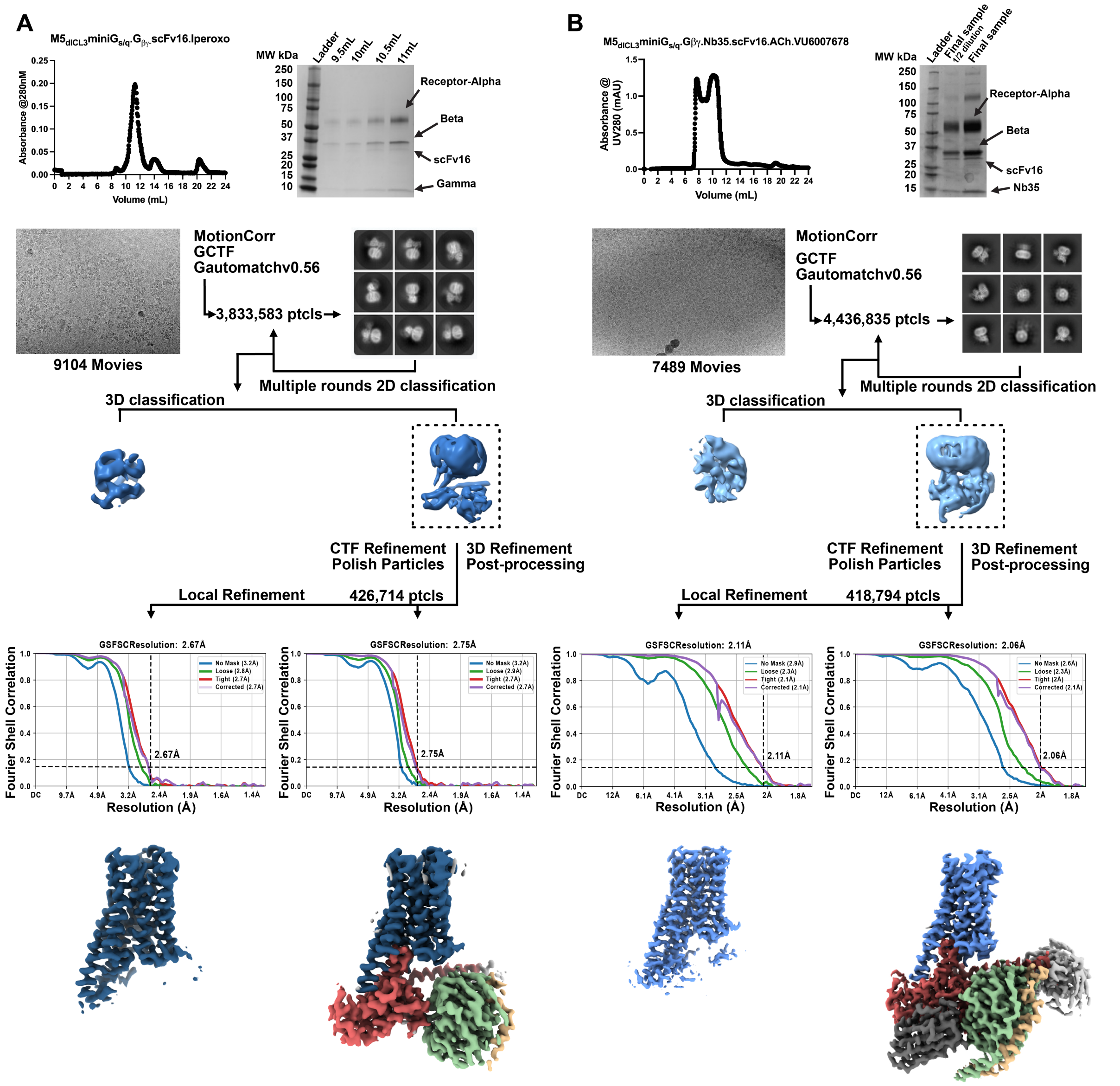

**Supplementary Figure 2. Cryo-EM data processing for the Iperoxo-bound $M_5$ mAChR-mini-G$\alpha_{qiN}$ and ACh-VU6007678-bound M5 mAChR-mini-G$\alpha_{qiN}$ complexes.** Flow chart for cryo-EM analysis as described in methods leading to final cryo-EM density maps for **(A)** the Iperoxo-bound $M_5$ mAChR-mini-G$\alpha_{mGsQi}$ complex and **(B)** ACh-VU6007678-bound $M_5$ mAChR-mini-G$\alpha_{mGsQi}$ complex.

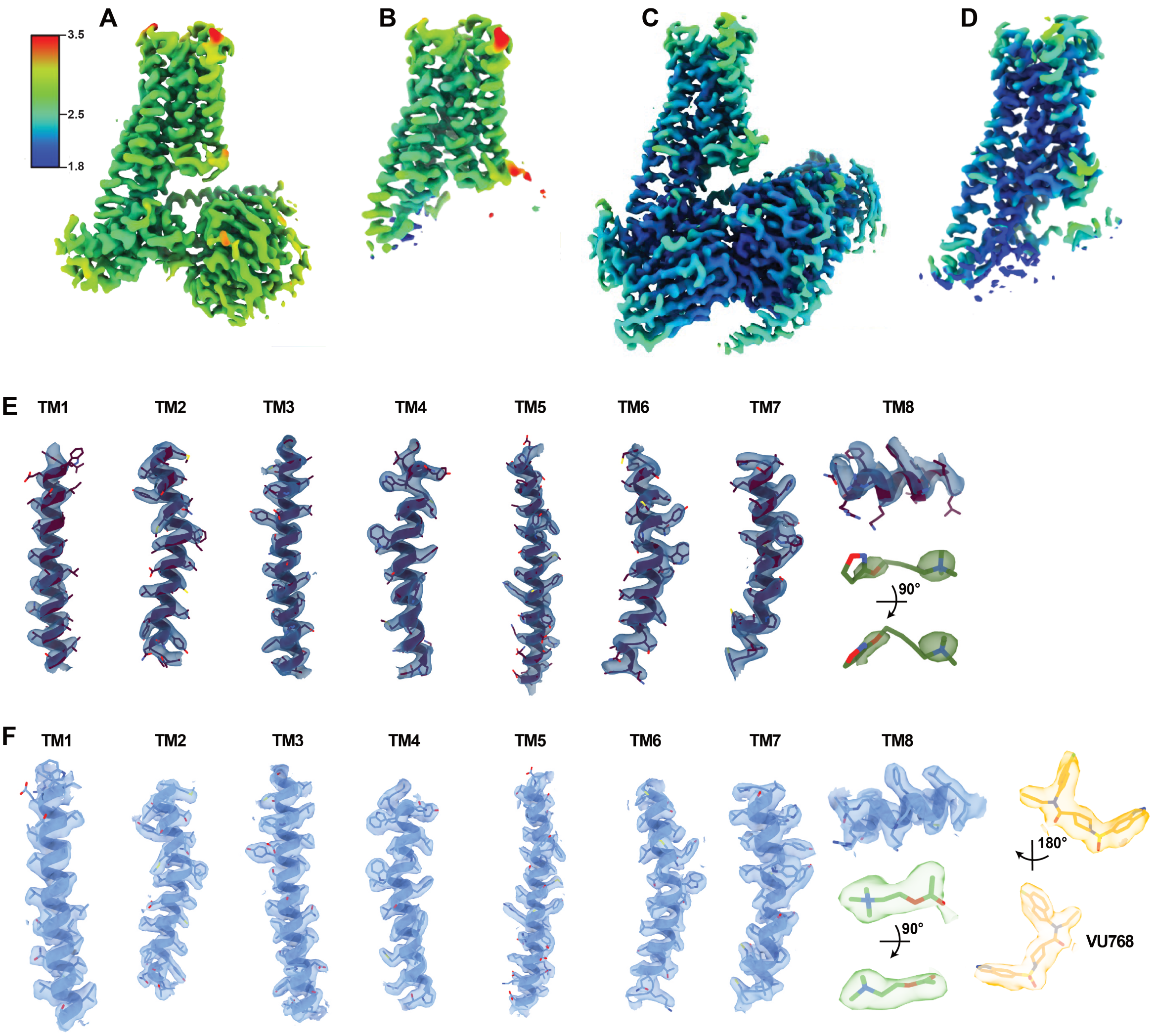


**Supplementary Figure 3. Cryo-EM density maps.** Local resolution map of the **(A)** complex, **(B)** receptor maps for the Iperoxo-bound $M_5$ mAChR-mini-G$\alpha_{mGsQi}$ structure and **(C)** complex, **(D)** receptor maps for the ACh-VU6007678-bound $M_5$ mAChR-mini-G$\alpha_{mGsQi}$ structure calculated in RELION 3.1. **(E-F)** Shown as cartoons the transmembrane domains of the $M_5$ mAChR and the α5 of $G_q$ modeled into the receptor focused cryo-EM maps, which is shown as a transparent surface contoured at 0.332 for (E) and 0.344 for (F).

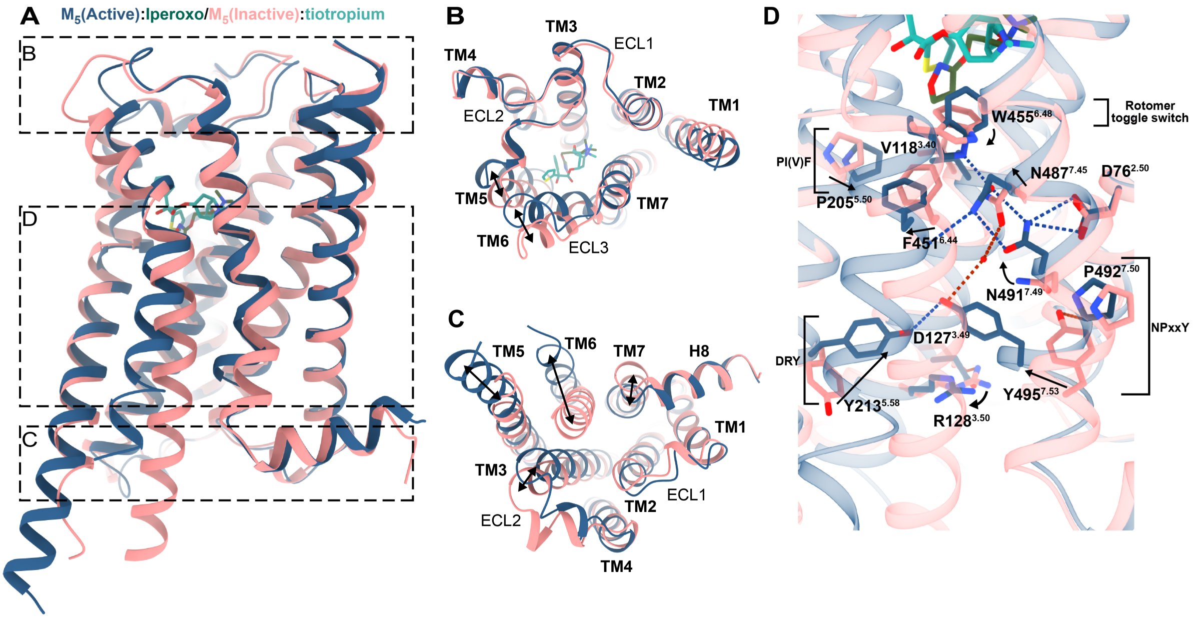


**Supplementary Figure 4. Comparison of active and inactive $M_5$ mAChR. (A)** Overlay of active, iperoxo bound $M_5$ mAChR (shown in dark blue) with inactive, tiotropium bound $M_5$ mAChR (shown in salmon, PDB: 6OL9). **(B)** Extracellular view comparing ECLs and TM regions. Arrows indicate movement between inactive and active state. **(C)** Intracellular view (G protein removed) comparing ICLs and TM regions. Arrows indicate movement between inactive and active state. **(D)** Key receptor activation motifs for class A GPCRs. Arrows indicate movement of residues between inactive and active state. Red dashed lines indicate hydrogen bonds in the inactive structure. Blue dashed lines indicate hydrogen bonds in the active structure.

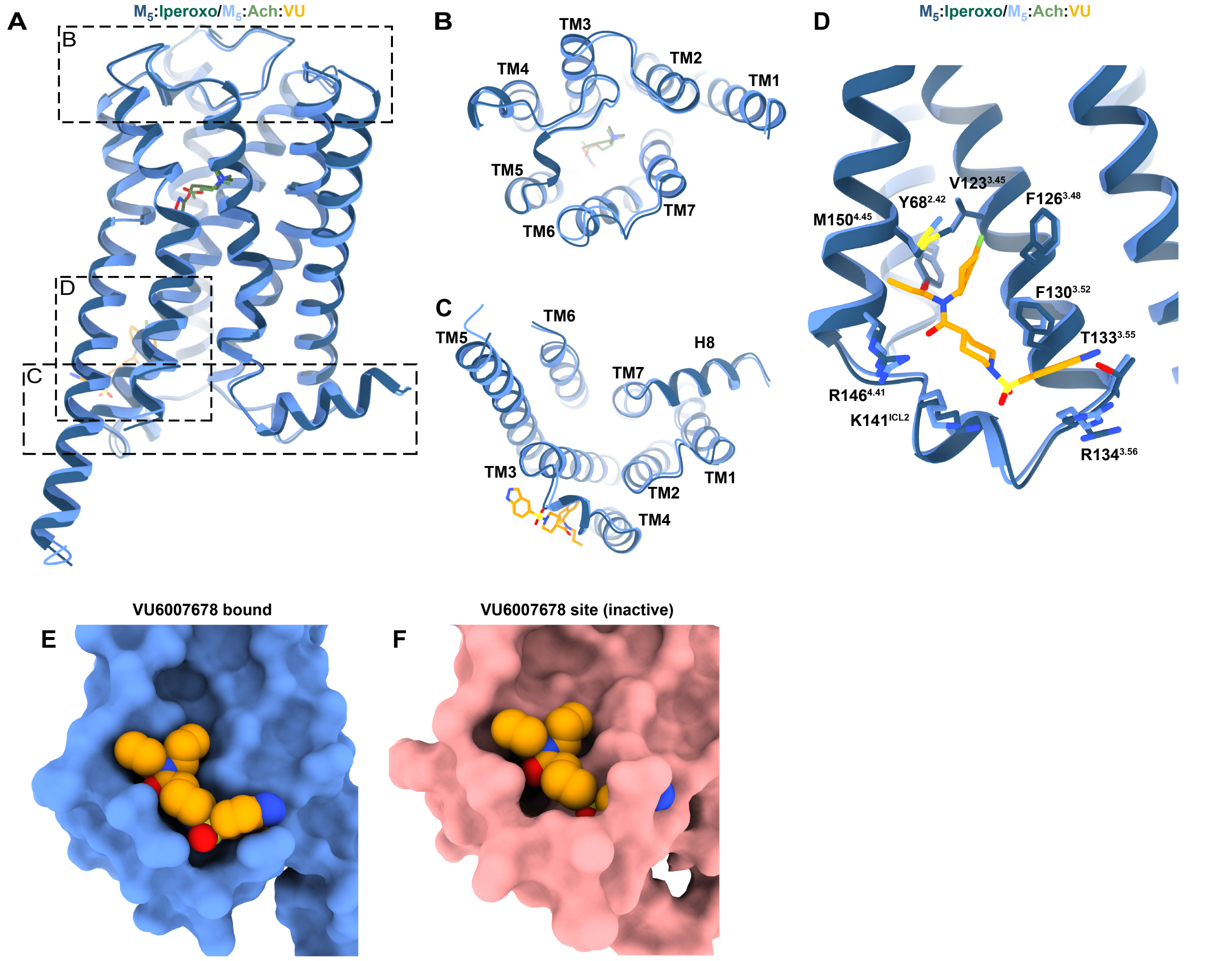


**Supplementary Figure 5. Comparison of iperoxo bound to ACh-VU6007678 bound M$_5$ mAChR. (A)** Overlay of iperoxo bound M$_5$ mAChR (shown in dark blue) with ACh-VU6007678 bound M$_5$ mAChR (shown in light blue). Iperoxo is shown in dark green sticks, ACh in light green sticks and VU6007678 in orange sticks. **(B)** Extracellular view comparing ECLs and TM regions. **(C)** Intracellular view (G protein removed) comparing ICLs and TM regions. **(D)** Comparison of the allosteric binding site at the ACh-VU6007678 bound M$_5$ mAChR to the iperoxo bound M$_5$ mAChR. Key residues are shown as sticks, iperoxo bound M$_5$ mAChR in dark blue, ACh-VU6007678 M$_5$ mAChR bound in light blue, VU6007678 is represented as orange sticks. **(E-F)** Spherical representation of the VU6007678 allosteric binding site at (E) ACh-VU6007678 bound M$_5$ mAChR and (F) inactive, tiotropium bound M$_5$ mAChR (PDB:6OL9).

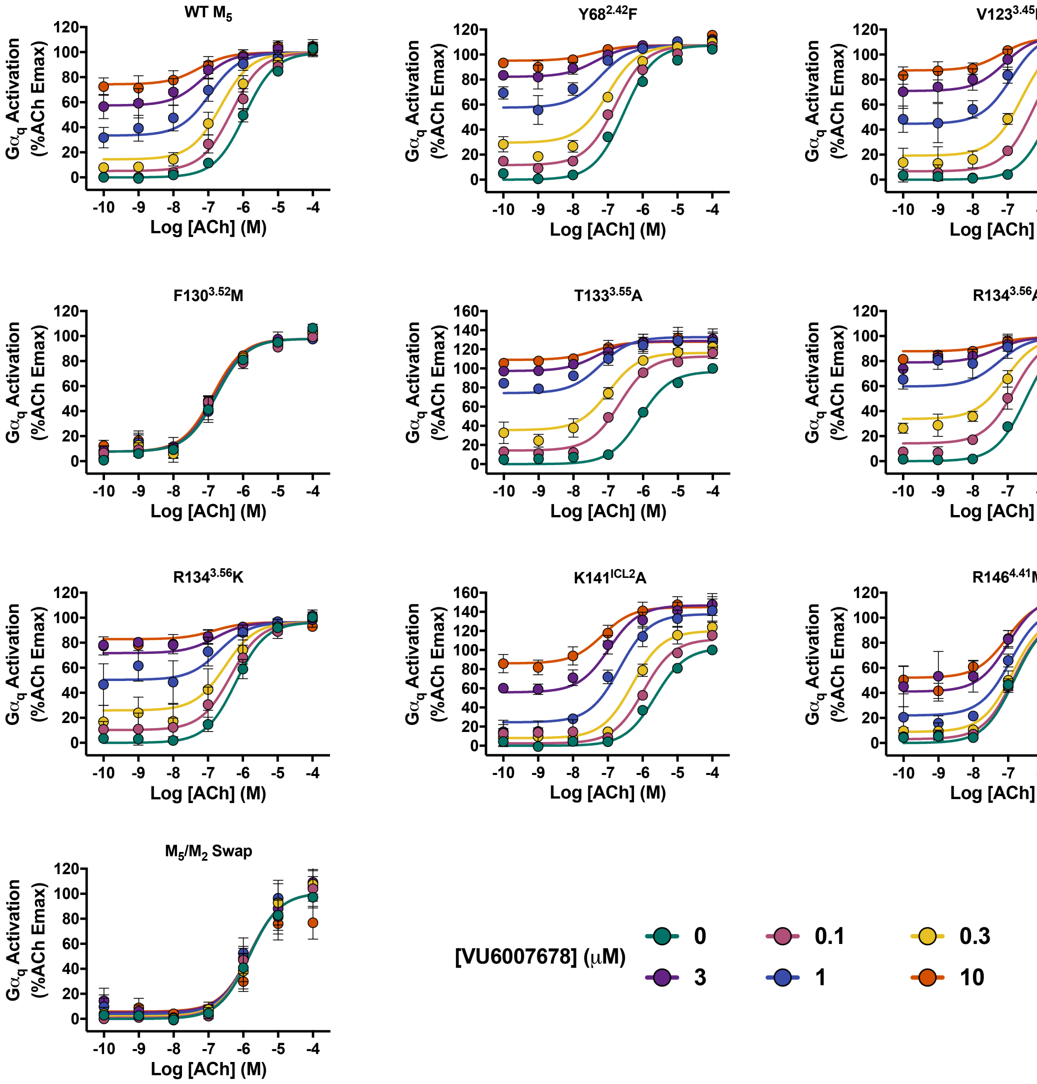


**Supplementary Figure 6. Functional interaction of VU6007678 with ACh at allosteric site mutants.** Interaction of VU6007678 with ACh in a Trupath $G\alpha_q$ activation assay at allosteric site mutant $M_5$ mAChRs expressed in CHO cells. Data points represent mean ± SEM of three to eight individual experiments performed in duplicate. WT $M_5$ mAChR *n* = 8, $M_5/M_2$ swap, $F130^{3.52}M$, $T133^{3.55}A$, $R134^{3.56}A$, $R134^{3.56}K$ mutants *n* = 3, $Y68^{2.42}F$, $V123^{3.45}I$, $K141^{34.56}A$, $R146^{4.41}M$ mutants *n* = 4. An operational model of allosterism was fit to the data. Parameters obtained are listed in Supplementary Table 4. Graphs for WT $M_5$ mAChR and $M_5/M_2$ swap were reproduced from main Figure 5B-C.

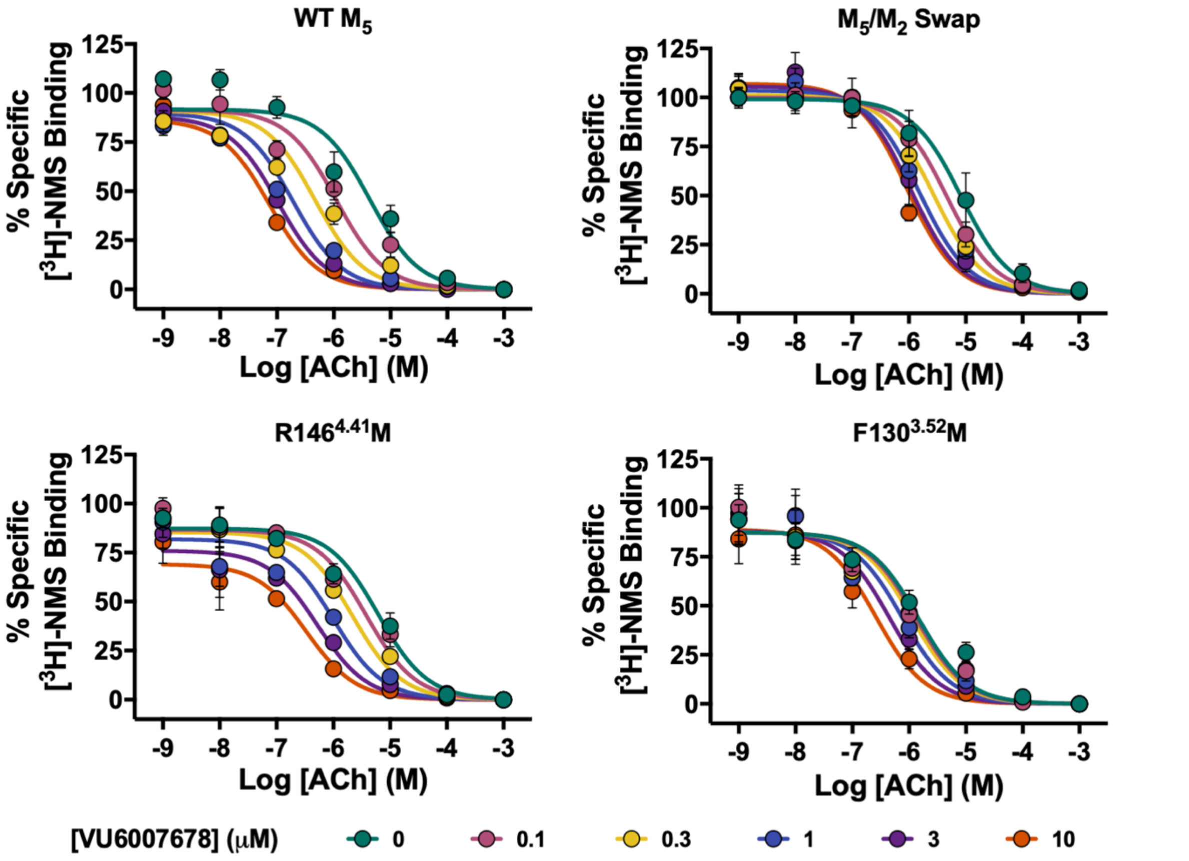


**Supplementary Figure 7. Binding interaction of VU6007678 with ACh at allosteric site mutants.** [$^3$H]-NMS equilibrium radioligand binding studies between ACh, VU6007678 and [$^3$H]-NMS. Data points represent mean ± SEM of three to five individual experiments performed in duplicate. WT $M_5$ mAChR *n* = 3, F130$^{3.52}$M, R146$^{4.41}$M *n* = 4, $M_5/M_2$ swap *n* = 5. Data was fit to an allosteric ternary complex model. Parameters obtained listed in Supplementary Table 4.

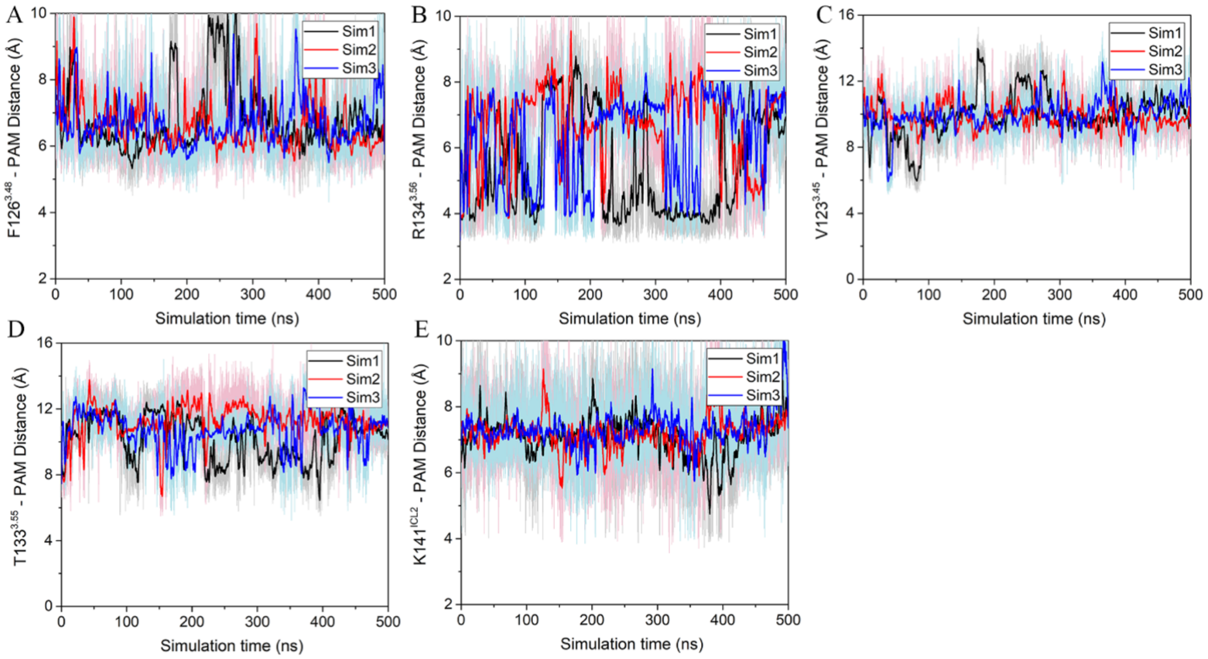


**Supplementary Figure 8.** Time courses of distances from VU6007678 to allosteric binding site residues in Å calculated from Gaussian accelerated molecular dynamics (GaMD) simulations performed with three separate replicates as indicated through different coloured traces.

**Supplementary Table 1 | Functional parameters for the ACh and ML380 interaction in IP1 accumulation assays.**

| Mutant | IP1 accumulation for the interaction of ML380 vs. ACh | | | |
|---|---|---|---|---|
| | ACh $pEC_{50}$[a] | $pK_B$[b] | Log $\tau_B$[c] | Log $\alpha\beta$[d] |
| **WT** | 6.94 ± 0.08 (7) | 5.43 ± 0.14 (7) | 0.32 ± 0.09 (7) | 1.48 ± 0.16 (7) |
| **EH4 Pocket** | 6.12 ± 0.11* (3) | 4.80 ± 0.38 (3) | 0.64 ± 0.30 (3) | 1.74 ± 0.39 (3) |
| **Y87$^{2.61}$A** | 6.40 ± 0.10* (3) | 6.34 ± 0.21* (3) | -0.85 ± 0.19* (3) | 0.83 ± 0.11* (3) |
| **Y90$^{2.64}$A** | 4.87 ± 0.06* (3) | 5.26 ± 0.11 (3) | -1.35 ± 0.37* (3) | 1.86 ± 0.09 (3) |
| **I91$^{2.65}$A** | 6.79 ± 0.10 (3) | 5.73 ± 0.21 (3) | -0.09 ± 0.08 (3) | 0.91 ± 0.25* (3) |
| **E182$^{ECL2}$A** | 6.94 ± 0.09 (3) | 5.95 ± 0.24 (3) | 0.12 ± 0.07 (3) | 0.69 ± 0.33* (3) |
| **Q184$^{ECL2}$A** | 6.48 ± 0.13* (3) | 5.42 ± 0.19 (3) | 0.02 ± 0.11 (3) | 1.69 ± 0.22 (3) |
| **S465$^{6.58}$A** | 6.90 ± 0.18 (3) | 5.54 ± 0.31 (3) | 0.12 ± 0.15 (3) | 1.38 ± 0.37 (3) |
| **D469$^{6.62}$A** | 6.71 ± 0.12 (3) | 5.59 ± 0.21 (3) | 0.15 ± 0.10 (3) | 1.64 ± 0.27 (3) |
| **K470$^{ECL3}$A** | 6.66 ± 0.10 (3) | 5.52 ± 0.16 (3) | -0.01 ± 0.08 (3) | 1.31 ± 0.19 (3) |
| **V474$^{7.32}$A** | 6.44 ± 0.12* (3) | 5.37 ± 0.22 (3) | -0.17 ± 0.10 (3) | 1.30 ± 0.22 (3) |
| **W477$^{7.35}$A** | 5.15 ± 0.07* (3) | 5.51 ± 0.12 (3) | -1.14 ± 0.26* (3) | 1.52 ± 0.09 (3) |
| **H478$^{7.36}$A** | 6.42 ± 0.12* (3) | 5.68 ± 0.12 (3) | -0.21 ± 0.10 (3) | 1.44 ± 0.19 (3) |

Data represent the mean ± S.E.M. of (n) independent experiments performed in duplicate. *, significantly different from WT, $p < 0.05$, one-way ANOVA, Dunnett's post hoc test.

[a] Negative logarithm of the concentration of ACh required to give half maximal response.

[b] Negative logarithm of the allosteric modulator equilibrium dissociation constant.

[c] Logarithm of operational efficacy parameter.

[d] Logarithm of functional cooperativity between agonist and allosteric modulator.

**Supplementary Table 2 | Data collection and refinement statistics**

| **Data Collection** | **$M_{5\Delta ICL3}$mAChR/m$G_{qNi}$/scFv16/Iperoxo** | **$M_{5\Delta ICL3}$mAChR/m$G_{qNi}$/scFv16/Nb35/ACh/VU6007678** |
|---|---|---|
| **PDB/EMDB** | 9EK0 / EMD-48111 | 9EJZ / EMD-48110 |
| **Micrographs** | 9104 | 7489 |
| **Electron dose ($e^-$/Å$^2$)** | 10.57 | 10.57 |
| **Voltage (kV)** | 300 | 300 |
| **Pixel size (Å)** | 0.65 | 0.82 |
| **Defocus range (μM)** | 0.5 - 1.5 | 0.5 - 1.5 |
| **Symmetry Imposed** | C1 | C1 |
| **Particles (final map)** | 426,714 | 418,794 |
| **Resolution (0.143 FSC) (Å)** | 2.75 | 2.06 |
| **Refinement** | | |
| **CCmap_model** | 0.84 | 0.88 |
| **Model Quality** | | |
| **R.m.s deviations** | | |
| **Bond Length (Å)** | 0.004 | 0.005 |
| **Bond angles (°)** | 0.722 | 0.728 |
| **Ramachandran** | | |
| **Favoured (%)** | 98.17 | 98.24 |
| **Outliers (%)** | 0 | 0 |
| **Rotamer outliers (%)** | 0.13 | 0.09 |
| **C-Beta deviations (%)** | 0 | 0 |
| **Clashscore** | 4.19 | 5.15 |
| **MolProbity score** | 1.20 | 1.27 |

**Supplementary Table 3 | Binding parameters for WT M5, M5-M2 TM chimeras**

| | Saturation Binding with [$^3$H]-NMS | | Interaction binding between [$^3$H]-NMS and ACh in the presence of ML380 | | | |
|---|---|---|---|---|---|---|
| | p$K_D$ ([$^3$H]-NMS)[a] | Sites per cell[b] | p$K_I$ (ACh)[c] | Log α (ACh)[d] | p$K_B$ (ML380)[e] | Log α ([$^3$H]-NMS)[f] |
| **WT M5** | 9.12 ± 0.14 (4) | 1363757 ± 198445 (4) | 5.04 ± 0.07 (3) | 0.84 ± 0.18 (3) | 5.40 ± 0.20 (3) | -0.47 ± 0.13 (3) |
| **M5 – M2 TM2, 3, 4** | 9.32 ± 0.16 (4) | 486397 ± 137869 (4) | 4.65 ± 0.08 (3)* | N.R. | N.R. | N.R. |
| **M5 – M2 TM3, 4, 5** | 9.23 ± 0.27 (3) | 1328988 ± 258858 (3) | 4.10 ± 0.07 (3)* | N.R. | N.R. | N.R. |
| **M5 – M2 TM1, 7, h8** | 8.34 ± 0.03 (4) | 443589 ± 53179 (4) | 4.75 ± 0.11 (3)* | 2.49 ± 0.55 (3) | 3.84 ± 0.51 (3) | =0[g] |

Data represent the mean ± S.E.M. of (n) independent experiments performed in duplicate. *, significantly different from WT, one-way ANOVA, Dunnett's post-hoc test, ($P$ < 0.05). N.D. not determined. N.R. no response

[a] Negative logarithm of the radioligand ([$^3$H]-NMS) equilibrium dissociation constant.

[b] Number of [$^3$H]-NMS binding sites per cell.

[c] Negative logarithm of the orthosteric agonist (ACh) equilibrium dissociation constant obtained from the allosteric ternary complex model.

[d] Logarithm of affinity cooperativity between the orthosteric agonist (ACh) and allosteric modulator (ML380).

[e] Negative logarithm of the allosteric modulator (ML380) equilibrium dissociation constant.

[f] Logarithm of affinity cooperativity between the orthosteric radioligand ([$^3$H]-NMS) and allosteric modulator (ML380).

[g] Value was not significantly different to zero and such was constrained to this value.

**Supplementary Table 4 | Functional and binding parameters for the ACh and VU6007678 interaction in TruPath G protein activation and [$^3$H]-NMS radioligand binding assays.**

| | TruPath G protein activation | | | | [$^3$H]-NMS radioligand binding | | | | |
|---|---|---|---|---|---|---|---|---|---|
| **Mutant** | **ACh pEC$_{50}$**$^a$ | **p$K_B$**$^b$ | **log $\tau_B$**$^c$ | **log αβ**$^d$ | **p$K_D$**$^e$ | **Sites per cell**$^f$ | **p$K_B$**$^g$ | **log α (ACh)**$^h$ | **log α ([$^3$H]-NMS)**$^h$ |
| **WT** | 5.95 ± 0.09 (8) | 4.94 ± 0.24 (8) | 0.80 ± 0.2 (8) | 2.24 ± 0.27 (8) | 9.36 ± 0.08 (5) | 587164 ± 66516 (5) | 5.67 ± 0.18 (3) | 1.83 ± 0.17 (3) | -0.06 ± 0.05 (3) |
| **Y68$^{2.42}$F** | 6.53 ± 0.06 (4)* | 4.76 ± 0.28 (4) | 1.29 ± 0.28 (4) | 2.24 ± 0.30 (4) | 9.30 ± 0.09 (4) | 641930 ± 103717 (4) | N.D. | N.D. | N.D. |
| **V123$^{3.45}$I** | 5.80 ± 0.12 (4) | 5.02 ± 0.26 (4) | 0.56 ± 0.22 (4) | 2.31 ± 0.29 (4) | 9.18 ± 0.09 (3) | 1054369 ± 31893 (3)* | N.D. | N.D. | N.D. |
| **F130$^{3.52}$M** | 6.75 ± 0.09 (3)* | N.D. | N.D. | N.D. | 9.40 ± 0.13 (4) | 782078 ± 98311 (4) | 5.07 ± 0.69 (4) | 0.99 ± 0.41 (4) | 0.03 ± 0.12 (4) |
| **T133$^{3.55}$A** | 6.09 ± 0.11 (3) | 5.19 ± 0.21 (3) | 1.12 ± 0.20 (3) | 2.35 ± 0.25 (3) | 9.18 ± 0.05 (4) | 419277 ± 63140 (4) | N.D. | N.D. | N.D. |
| **R134$^{3.56}$A** | 6.48 ± 0.09 (3)* | 5.08 ± 0.27 (3) | 0.97 ± 0.26 (3) | 2.11 ± 0.31 (3) | 9.27 ± 0.27 (4) | 881469 ± 131189 (4) | N.D. | N.D. | N.D. |
| **R134$^{3.56}$K** | 6.20 ± 0.11 (3) | 4.98 ± 0.36 (3) | 1.06 ± 0.33 (3) | 1.78 ± 0.43 (3) | 9.44 ± 0.08 (3) | 629606 ± 90141 (3) | N.D. | N.D. | N.D. |
| **K141$^{ICL2}$A** | 5.64 ± 0.14 (4) | 4.75 ± 0.30 (4) | 0.62 ± 0.25 (4) | 2.34 ± 0.32 (4) | 9.28 ± 0.09 (4) | 566941 ± 93126 (4) | N.D. | N.D. | N.D. |
| **R146$^{4.41}$M** | 6.83 ± 0.07 (4)* | 5.47 ± 0.17 (4) | -0.14 ± 0.11 (4)* | 0.57 ± 0.16 (4)* | 9.31 ± 0.21 (4) | 897976 ± 92116 (4) | 5.54 ± 0.23 (4) | 1.30 ± 0.20 (4) | -0.23 ± 0.06 (4) |
| **M$_5$-M$_2$ Swap** | 5.82 ± 0.13 (3) | N.D. | N.D. | N.D. | 9.31 ± 0.16 (4) | 466347 ± 98510 (4) | 5.98 ± 0.24 (5) | 1.02 ± 0.14 (5) | 0.07 ± 0.04 (5) |

Data represent the mean ± S.E.M. of (n) independent experiments performed in duplicate. *, significantly different from WT, $p < 0.05$, one-way ANOVA, Dunnett's post hoc test. N.D., not determined.

$^a$ Negative logarithm of the concentration of ACh required to give half maximal response obtained from the three-parameter logistic equation.

$^b$ Negative logarithm of the allosteric modulator (VU6007678) equilibrium dissociation constant (p$K_B$) obtained from the operational model of allosterism.

$^c$ Logarithm of operational efficacy parameter obtained from the operational model of allosterism corrected for receptor expression (sites per cell).

$^d$ Logarithm of functional cooperativity between agonist (ACh) and allosteric modulator (VU6007678) obtained from the operational model of allosterism.

$^e$ Negative logarithm of the radioligand ([$^3$H]-NMS) equilibrium dissociation constant.

$^f$ Number of [$^3$H]-NMS binding sites per cell.

$^g$ Negative logarithm of the allosteric modulator (VU6007678) equilibrium dissociation constant obtained from the allosteric ternary complex model.

$^h$ Logarithm of affinity cooperativity between the orthosteric agonist (ACh) or orthosteric radioligand ([$^3$H]-NMS) and allosteric modulator (VU6007678) obtained from the allosteric ternary complex model.
